# Host kinase regulation of *Plasmodium vivax* dormant and replicating liver stages

**DOI:** 10.1101/2023.11.13.566868

**Authors:** Elizabeth KK Glennon, Ling Wei, Wanlapa Roobsoong, Veronica I Primavera, Tinotenda Tongogara, Conrad B Yee, Jetsumon Sattabongkot, Alexis Kaushansky

## Abstract

Upon transmission to the liver, *Plasmodium vivax* parasites form replicating schizonts, which continue to initiate blood-stage infection, or dormant hypnozoites that reactivate weeks to months after initial infection. *P. vivax* phenotypes in the field vary significantly, including the ratio of schizonts to hypnozoites formed and the frequency and timing of relapse. Evidence suggests that both parasite genetics and environmental factors underly this heterogeneity. We previously demonstrated that data on the effect of a panel of kinase inhibitors with overlapping targets on *Plasmodium* liver stage infection, in combination with a computational approach called kinase regression (KiR), can be used to uncover novel host regulators of infection. Here, we applied KiR to evaluate the extent to which *P. vivax* liver-stage parasites are susceptible to changes in host kinase activity. We identified a role for a subset of host kinases in regulating schizont and hypnozoite infection and schizont size and characterized overlap as well as variability in host phosphosignaling dependencies between parasite forms and across multiple patient isolates. Striking, our data point to variability in host dependencies across *P. vivax* isolates, suggesting one possible origin of the heterogeneity observed across *P. vivax* in the field.

## Introduction

Malaria presents a significant global health burden with nearly 250 million clinical cases and more than a half million deaths annually world-wide ^1^. Although significant progress has been made towards reducing the incidence of malaria, the *Plasmodium* parasite has developed resistance to all drugs in widespread use ^2–4^. To initiate symptomatic blood-stage infection after transmission by a female *Anopheles* mosquito, the malaria parasite must first develop and replicate within the liver. Upon invading a hepatocyte, *Plasmodium* sporozoites develop as schizonts, growing and replicating to produce merozoites, which go on to invade red blood cells. A major hurdle to malaria eradication is the ability of *Plasmodium vivax* to form liver-resident dormant stages called hypnozoites in addition to the rapidly replicating schizont form. Unlike schizonts, hypnozoites do not initially replicate, and instead reactivate weeks to months after the initial infection.

Although no DNA replication occurs, hypnozoites are not quiescent. Hypnozoites exhibit a modest increase in size over the course of infection both *in vitro* and *in vivo* and exhibit changes in gene expression, membrane protein distribution, and exhibit altered sensitivities to schizonticidal drugs over time ^5–7^. Transcriptomic studies of *P. vivax* and *Plasmodium cynomolgi* dormant forms revealed enrichment of fatty acid synthesis and redox metabolism ^6,8,9^ suggesting that, despite their overall reduced transcriptional activity, hypnozoites are actively maintaining their redox state and may be susceptible to its perturbation.

It is likely that a combination of parasite genetics and the host environment influences both maintenance and reactivation of *P. vivax* liver-stage (LS) parasites. Given the potential for diversity in both parasite-intrinsic and environmental stimuli, it is not surprising that the phenotypes observed for *P. vivax* in the field are heterogeneous. The timing of relapse due to hypnozoite activation has been shown to exhibit both periodicity and random constant rate patterns, suggesting that there are both triggered and stochastic processes that mediate the conversion between dormant and activated states ^10^. Epidemiological studies suggest the rate of relapse varies ecogeographically ^11,12^, while others have demonstrated that inflammation, particularly blood-stage malaria, can trigger hypnozoite reactivation ^13–16^. More recent models of *P. vivax* recurrence support roles for population and temporal heterogeneity in determining overall relapse patterns ^17^. *P. vivax* isolates are traditionally divided into two variants, VK247 and VK210, based the amino acid sequence of circumsporozoite protein (CSP). These variants differ in their mosquito vectors and have been shown to produce different ratios of hypnozoites to schizonts in a humanized mouse model ^5,18^. However, the genetic diversity of *P. vivax* populations is not fully captured by this basic genotyping^19^. One longitudinal study identified 14 different *Pv*CSP alleles in patients at the Thai-Mayanmar border ^20^ while another identified 67 unique *P. vivax* merozoite surface protein 1 (*Pv*MSP-1) haplotypes ^21^. A large proportion of recurring *P. vivax* cases are heterologous infections and contain different parasite genotypes than acute infection ^22^, with a particularly high level of genetic diversity in parasites circulating in and around Thailand ^22,23^. *In vitro* work has also demonstrated a role for both parasite genetics and host environmental factors in regulating *P. vivax* infection, showing variation in infection rate, hypnozoite:schizont ratio, and schizont size across both parasite isolates and primary hepatocyte donors ^24^.

Work on schizont LS infection with *P. falciparum, P. yoelii, and P. berghei* has demonstrated parasite dependence on a multitude of host cell processes including the blocking apoptosis ^25–28^, regulation of lipid peroxidation ^29^, nutrient uptake ^30,31^, and re-orientation of vesicular trafficking to the parasitophorous vacuole ^32–35^. However, it is very unusual to compare the reliance on host regulators of infection between *Plasmodium* species, so we are limited in our ability to extrapolate among them. Where direct comparisons among species have been conducted, variation, as well as similarities, in host factor dependencies has been uncovered ^28,36–38^. *P. vivax* hypnozoites and schizonts have differing susceptibilities to many parasite-targeted drugs, but it remains an open question whether these parasite forms have common dependency on host determinants of infection. Inhibition of the water and small molecule channel aquaporin 3, which reduced LS *Plasmodium berghei* ^31^, also inhibited *P. vivax* schizonts and hypnozoites ^7^. *P. vivax* schizonts and hypnozoites exhibited overlapping susceptibility to treatment with interferons (IFN), with both IFNγ and IFNα reducing hypnozoite and schizont infection *in vitro* ^39^. While several broad genetic screens have been conducted to identify host genes that influence LS infection with other, more tractable, *Plasmodium* species ^32,33,40^, the relative intractability of primary hepatocytes to genetic manipulation and the difficulty of producing *P. vivax* sporozoites, make it particularly challenging to conduct large genetic screens. Currently, the full extent to which *P. vivax* LS infection is susceptible to manipulation of the host environment remains an open question.

To more broadly probe the host dependencies of *P. vivax* LS infection we have adapted kinase regression (KiR), which utilizes the characterized and quantified polypharmacology of kinase inhibitors and machine learning to predict which of 300 human kinases play a role in a cellular phenotype. In KiR, a set of ∼30 commercially available compounds with known activities against each of ∼300 human kinases ^41^ are tested for their capacity to inhibit any quantitative phenotype. Using machine learning, these two data sets are combined to generate predictions of kinases that contribute most substantially to the given cellular outcome. This approach was first pioneered to identify regulators of cell migration ^42^, and then determinants of cancer susceptibility ^43–45^, Kaposi’s sarcoma-associated herpesvirus reactivation ^46^, and SARS-CoV-2 induced cytokine release ^47^. We have utilized KiR to predict both previously described and novel host kinase regulators of *P. yoelii* LS infection ^48^ as well as to identify kinases that regulate the blood-brain endothelial barrier ^49^. We hypothesized that the host environment plays a major role in regulating *P. vivax* LS biology and have utilized KiR to probe the extent to which hypnozoites and schizonts are dependent on host phosphosignaling across multiple parasite isolates.

## Results

### *P. vivax* isolates exhibit different infection phenotypes in primary human hepatocytes

To probe the role of host kinase signaling in regulating *P. vivax* liver-stage (LS) infection we utilized the 384-well primary human hepatocyte microculture system described previously ^50^. We infected primary human hepatocytes at an MOI of ∼0.5 with sporozoites derived from three separate clinical isolates, spread across two independent infections, using the same batch of primary hepatocytes. We termed the three clinical isolates A, B, and C; infections from isolates B and C were performed at the same time. Parasite isolates were collected in the Tak province in northwestern Thailand and were all the VK210 CSP variant, which is highly prevalent in Thailand ^51,52^. 60,000 sporozoites were isolated per mosquito for isolate B and 64,000 per mosquito for isolate C. We allowed the parasites to develop for eight days post-infection and quantified hypnozoites and schizonts by microscopy. Liver stage parasites were identified by circumferential *P. vivax* Upregulated in Infectious Sporozoites-4 (*Pv*UIS4) staining. Schizonts were distinguished from hypnozoites by additional morphological features; schizonts were defined as having multiple nuclear masses and a diameter of over 10μm, and hypnozoites were defined as having a diameter of under 8μm and displaying the characteristic UIS4 prominence (**Fig. 1a**) ^5,53^. Even in the absence of any treatment, parasites displayed significantly varying phenotypes between the three isolates tested. Infection rates were significantly lower in isolate C with hypnozoite levels below robustly quantifiable levels (**Fig. 1b-c**). Isolates A and B had comparable overall infection rates but significantly different ratios of hypnozoites to schizonts (**Fig. 1b-c**). Schizont size spanned a large range and was significantly higher in isolate C compared to the other two isolates (**Fig. 1d**). The variance of schizont size was also significantly higher in isolate C compared to isolate A.

**Figure 1.**
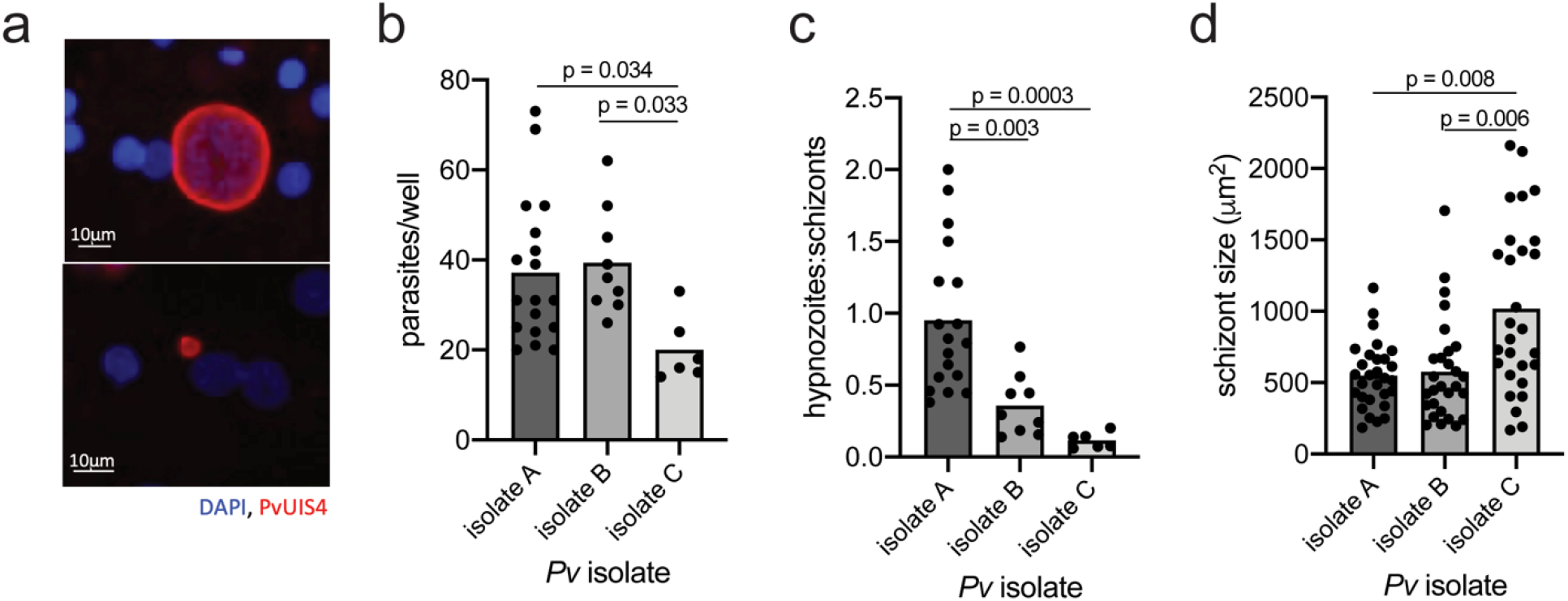
*P. vivax* parasites exhibit variation across different patient isolates. (a) Representative images of *P. vivax* schizonts (top) and hypnozoites (bottom) 8 dpi in primary human hepatocytes stained with DAPI and *Pv*<u>UIS4</u>. Comparison of (b) infection rate, (c) ratio of parasite forms, and (d) schizont size across three unique *P. vivax* isolates 8 dpi. Each dot represents counts from a control well. Data were analyzed by ANOVA with Tukey’s multiple comparisons test.

### Host kinase inhibition influences schizont and hypnozoite infection levels

We have previously demonstrated that, using a panel of 34 kinase inhibitors (KIs), novel kinase regulators of *Plasmodium* LS infection can be uncovered by KiR ^48^. To apply this to *P. vivax* infection, we infected primary hepatocytes with each of the three isolates and treated each infected culture with the panel of 34 inhibitors (**Supplementary Table 1**). Inhibitors previously shown to reduce *P. falciparum* blood-stage (BS) infection ^48^ were excluded from the screen, as we reasoned that they are more likely to be targeting *Plasmodium* kinases directly, making their activity difficult to interpret. One inhibitor, staurosporine (KI29), that exhibited significant toxicity over 20% in primary hepatocytes (**Supplementary Fig. 1**), was also removed from the screen. No other inhibitors exhibited statistically significant toxicity >20% across all three experiments. Schizonts and hypnozoites were quantified in response to treatment with the panel of inhibitors, with PI4 kinase inhibitor MMV390048 (PI4Ki) used as a positive schizont-targeting control in two of the three isolates (**Fig. 2**). Edge effects were tested for by comparing schizont and hypnozoite levels in technical replicates in outer edge or inner wells for each biological replicate. No significant effect of well position on infection was observed between paired wells that received the same treatment (**Supplementary Fig. 2**). PI4Ki dramatically reduced schizont infection as expected (data published in ^54^) while kinase inhibitors exhibited a range of effects across forms and isolates. Strikingly, many inhibitors exhibited different effects across the different isolates. However, some inhibitors exhibited a similar effect across isolates. Specifically, two inhibitors, GSK-3 inhibitor X (KI15) and K252a (KI20) consistently reduced and increased infection, respectively, for both schizonts and hypnozoites across all isolates, using a 20% change cutoff. Two additional KIs (KI31 and KI18) consistently reduced schizont infection across all isolates. Four additional KIs (KI6, KI8, KI10, KI11) consistently reduced, and two additional KIs (KI17 and KI30) consistently increased, hypnozoite infection levels.

**Figure 2.**
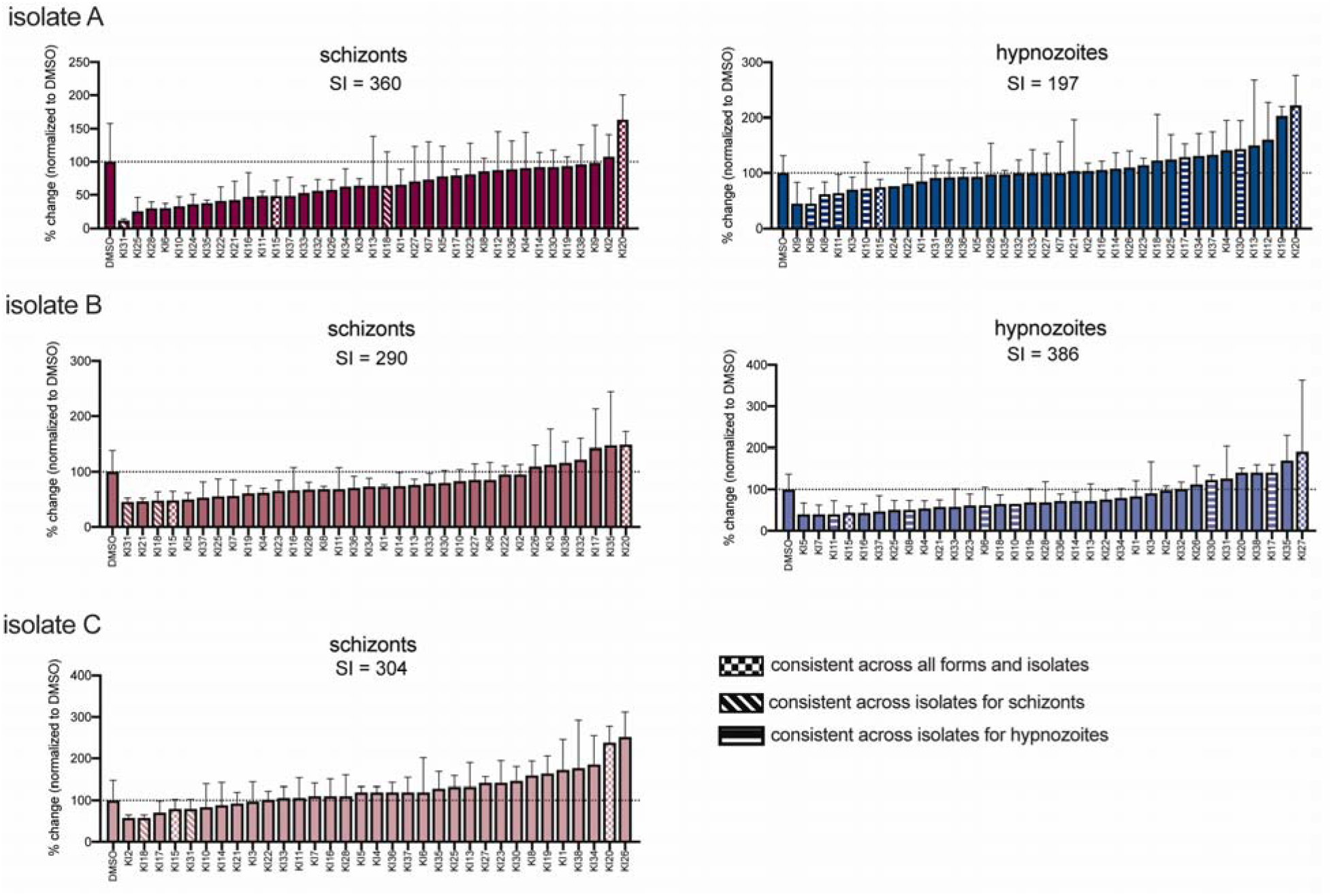
Kinase inhibitors alter infection rates with *P. vivax* schizonts and hypnozoites. Effect of panel of 28 kinase inhibitors on infection levels of three different *P. vivax* isolates in the same lot of primary human hepatocytes. Schizonts and hypnozoites were quantified 8 dpi from 3 technical replicates for each inhibitor and normalized to a DMSO control. Data are shown as mean and standard deviation of 3 technical replicates for each treatment. Inhibitors that consistently increased or decreased infection across forms and/or isolates, using a 20% change cut-off, are indicated. A sensitivity index (SI) was calculated for each form and isolate.

We next asked if schizonts were more susceptible to kinase inhibitor treatment, overall, than hypnozoites. However, a strict average of inhibition does not provide a robust measure of overall sensitivity to host phosphosignaling inhibition because kinase inhibitors within the panel vary tremendously in their level of promiscuity. The number of kinases inhibited by a single KI by >30% ranged from 1 to 228 within the panel of 300 kinases for which we have data ^41^. Within the panel of KIs, more promiscuous inhibitors did not necessarily inhibit infection by a greater extent. Comparing the number of kinases inhibited by each individual inhibitor with its effect on schizont or hypnozoite infection, for isolate A, showed a moderate positive correlation across a range of inhibition cut-off levels from 20-80%, with the most promiscuous KI in the screen, K252a (KI20), inducing the largest increase in infection rate for both schizonts and hypnozoites (**Supplementary Fig. 3**). To more rigorously evaluate how the overall phosphosignaling landscape impacts schizont and hypnozoite levels, we developed a “sensitivity index” (SI), which accounts for the polypharmacology of each kinase inhibitor by assigning each inhibitor a weight based on the extent of inhibition across the 300 kinases in the dataset (**Supplementary Table 1**) ^41^. These weights were then integrated with the effect of each inhibitor on schizont or hypnozoite infection to calculate an overall measure of sensitivity to kinase inhibition for each phenotype. Using this statistic, we observed variation in sensitivity both between forms and parasite isolates, with the dormant hypnozoite form not always showing less sensitivity to manipulation of host signaling (**Fig. 2**)

The ratio of hypnozoites to schizonts formed upon infection is known to be influenced by parasite genetics ^11,12^. However, there is also evidence that the host environment can influence hypnozoite ratio ^24^ and reactivation ^13–16^. We asked if our data were consistent with the hypothesis that a reduction in schizonts or hypnozoites upon KI treatment was due to a shift in forms: more schizonts corresponding with fewer hypnozoites or vice versa. If this hypothesis was true, we would expect to observe an inverse correlation between the impact on schizont and hypnozoite numbers across treatments. In contrast, we observe a strong positive correlation between the impact of a given inhibitor on schizonts and hypnozoites, demonstrating that drugs that inhibit schizonts are more likely to also inhibit hypnozoites (**Fig. 3**). Using a 20% change cut-off, two KIs were identified in each of two isolates that increased hypnozoite infection while decreasing schizont infection, however which KIs had this effect was not conserved between isolates. No KIs decreased hypnozoite infection while increasing schizont infection.

**Figure 3.**
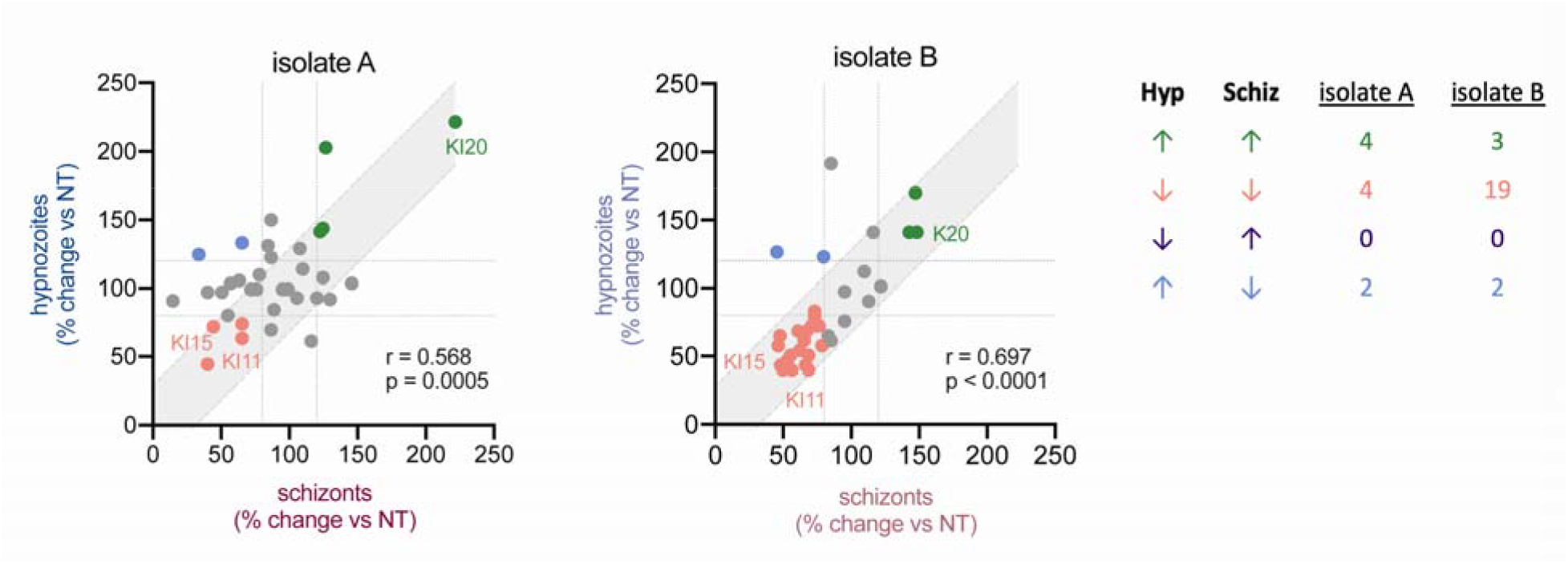
Effect of KIs on hypnozoites and schizonts correlates within each isolate. Effect of each inhibitor on schizont infection plotted against its effect on hypnozoite infection for each isolate. Each dot represents the mean % change compared to DMSO controls for a single kinase inhibitor. Gray shaded area indicates a 95% confidence interval, based on variation of control wells, around x=y. A 20% change cut-off, indicated by the dotted gray lines, was used to categorize the effect of inhibitors on each parasite form.

### Using kinase regression to predict susceptibility of *P. vivax* liver-stages to kinase inhibitors

Kinase regression (KiR) utilizes the overlapping specificities of a small panel of kinase inhibitors to deconvolve the effects of 300 host kinases, and over 100 additional kinase inhibitors not in the screen, on a quantifiable phenotype (**Fig. 4a**) ^42^. We previously used KiR to identify kinases and kinase inhibitors that altered *Plasmodium yoelii* infection, with 85% accuracy ^48^. Because each parasite isolate originated from an independent patient and the impact of inhibitors varied across isolates (**Fig. 2**) we used KiR to make predictions on each isolate individually (**Supplementary Table 2**). We first used the data from isolate A to predict the effect of 178 inhibitors on infection with each parasite form (**Fig. 4b**). We then tested the effect of five new KIs predicted to reduce schizont infection by > 80%, based on data from isolate A, on schizont infection in isolate B over a range of concentrations. Consistent with the variation in the response of different parasite isolates to kinase inhibition, two of the five inhibitors, SB203580 and JNK inhibitor V, significantly reduced schizont infection in isolate B at any of the concentrations tested (**Fig. 4c**). We also conducted dose response curves in isolate B for two inhibitors that were part of the original screen: SU11274 (KI31), which reduced schizont infection across isolates A and B, and casein kinase I inhibitor D4476 (KI6), which reduced infection in isolate A but not isolate B (**Supplementary Fig. 4a**). KI31 (SU11274) reduced schizont infection with increasing magnitude as concentration increased (**Supplementary Fig. 4b**). KI6 (casein kinase I inhibitor D4476), which did not significantly impact infection of isolate B in the screen (500nM), only reduce infection at the highest concentration tested (10μM) (**Supplementary Fig. 4b**).

**Figure 4.**
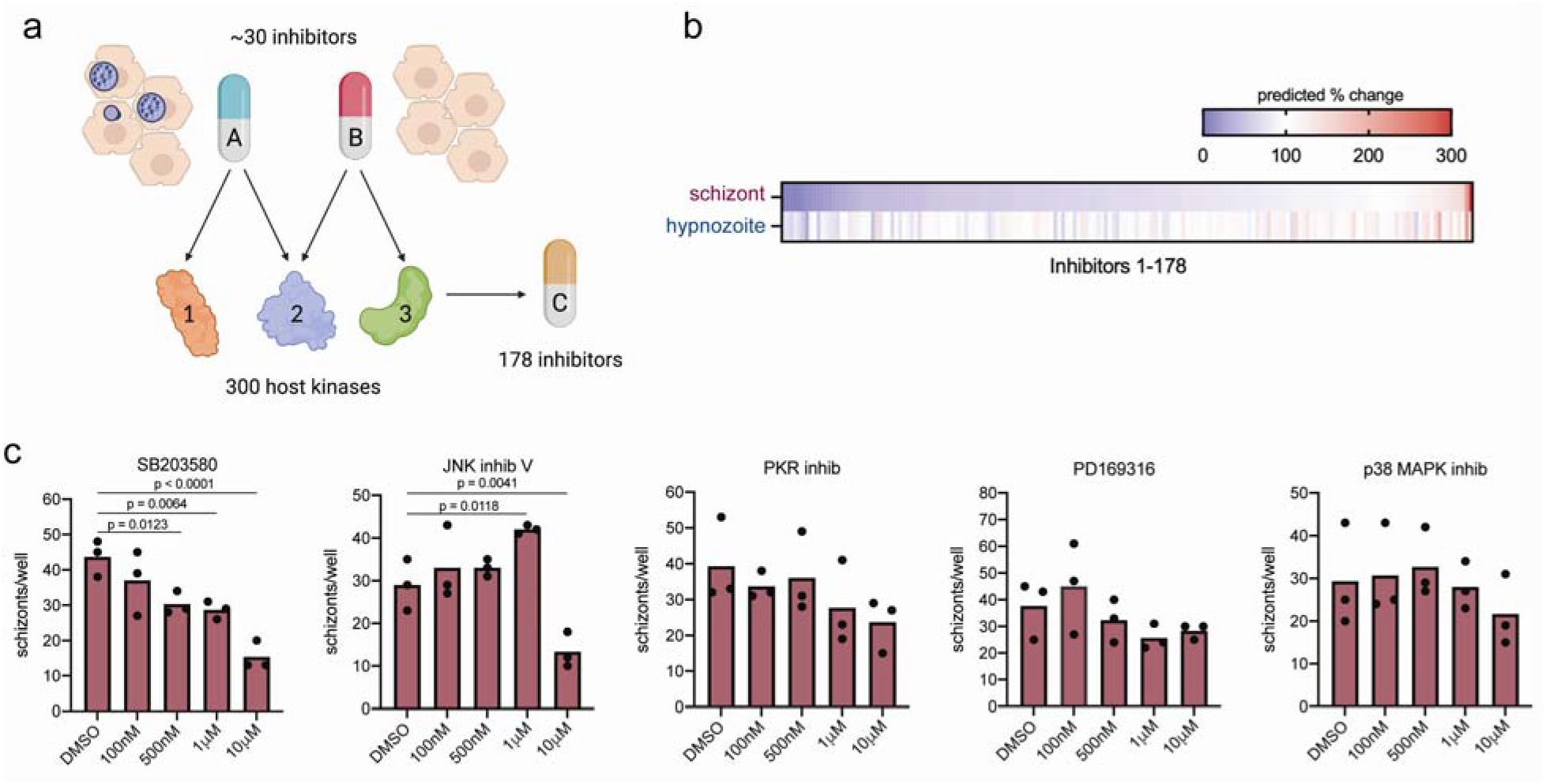
Kinase regression predicts kinase inhibitors that alter *P. vivax* infection. (a) Schematic of kinase regression. Briefly, the effect a panel of ∼30 inhibitors, with characterized overlapping kinase specificities, on LS infection was measured. These data were then used to predict the role of 300 host kinases on LS infection as well as the effect of 178 kinase inhibitors. (b) Heatmap showing predicted effect of 178 inhibitors on schizont infection based on data from isolate A (Fig. 2). (c) Schizont dose response curves in isolate B for five inhibitors that were predicted to reduce infection in isolate A. Primary human hepatocytes were infected with *P. vivax* sporozoites from isolate B, treated 24h post-invasion, and then fixed 8 dpi for parasite quantification. Each dot represents a technical replicate. Data were analyzed by Fisher’s LSD test.

To more broadly probe the predictive power between isolates for susceptibility to host kinase inhibition we used the KI screen data from one isolate to predict drug susceptibility and then compared those predictions to the KI screen data from another isolate (see Methods). Predictions and screen data showed only a weak correlation for schizont infection rates from separate isolates (**Fig. 5a**) and no correlation for hypnozoites from separate isolates (**Fig. 5b**). As we had observed some consistency in the effects of KIs on hypnozoites and schizonts within an isolate (**Fig. 2,3**) we asked if data from one form could predict activity for the other form within the same isolate. Kinase inhibitor susceptibility predictions made using screen data from schizonts correlated strongly with the drug screen data for hypnozoites within each isolate (**Fig. 5c**), suggesting isolate identity may play a stronger role than parasite stage in determining *P. vivax* LS susceptibility to host kinase perturbation.

**Figure 5.**
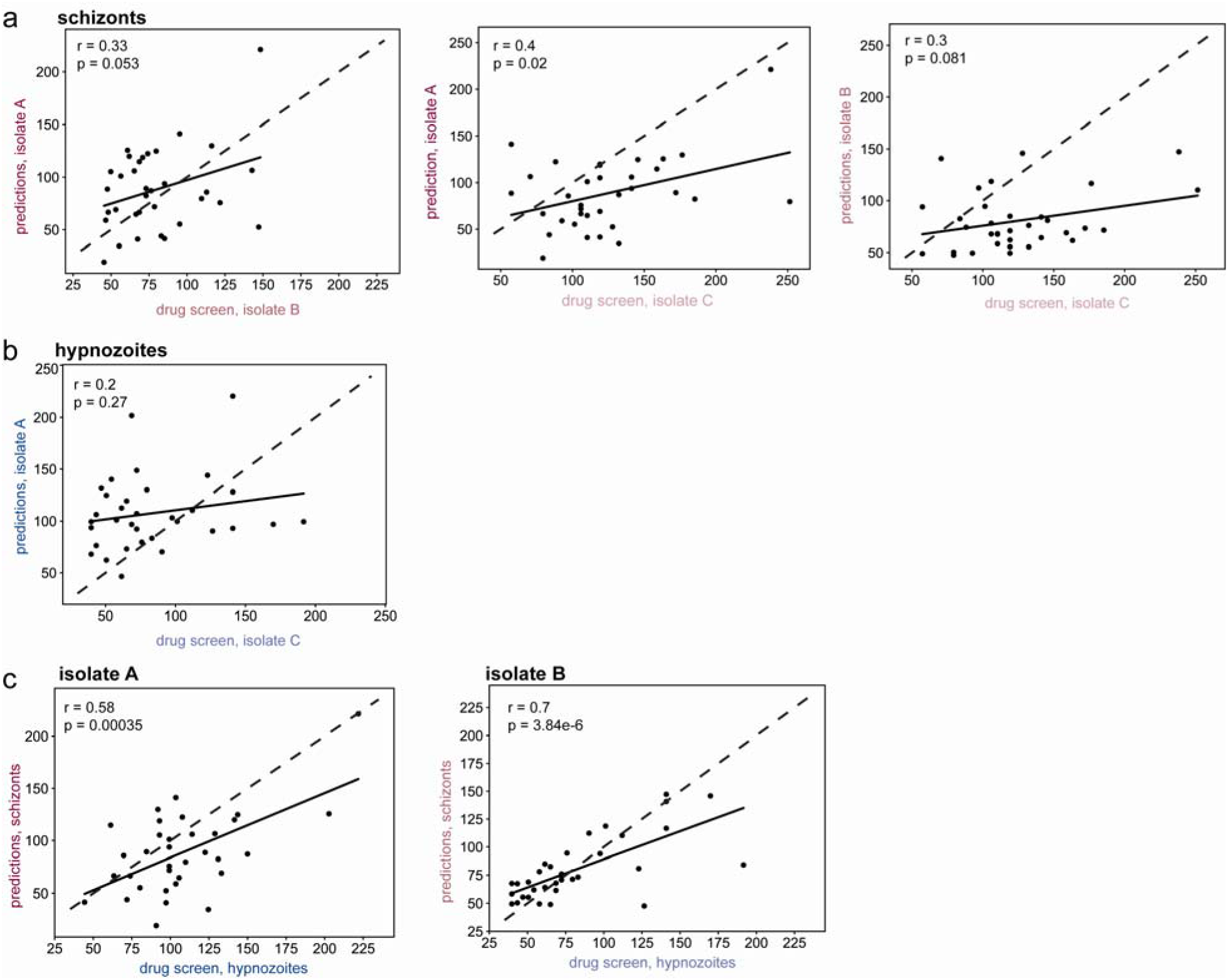
Predictive power of KiR across parasite isolates and forms. KI effect on (a) schizont or (b) hypnozoite infection was predicted using data from one isolate and graphed against drug screen data from another isolate. (c) KI effect on infection was predicted using schizont data and graphed against drug screen data for hypnozoites within the same parasite isolate. Data are shown as percent change compared to DMSO controls at 8 dpi. Each dot represents a single inhibitor. Pearson correlation coefficients (r) were calculated for each comparison.

### Host kinase inhibition regulates schizont size

Multiple lines of research have indicated that parasite-intrinsic and/or host factors might influence the rate of schizont growth. When comparing four hepatocyte donors, Vantaux and colleagues demonstrated that *P. vivax* LS schizonts can vary dramatically in size across different hepatocyte donors ^24^, although they observed reasonably modest differences in schizont size when comparing isolates. In contrast, we observed significant differences in both average and variance of schizont size between *P. vivax* isolates within the same hepatocyte donor (**Fig. 1**). Parasite size likely serves as a surrogate for parasite biomass, as in untreated wells schizont size correlated strongly with the parasite nuclear stain signal (**Supplementary Fig. 5a**).

When we measured the effect of the kinase inhibitor panel on the size of schizonts, we again saw minimal overlap in effect between isolates (**Fig. 6**). Two KIs consistently reduced schizont size, one of which (SU11274/KI31) had also consistently reduced schizont infection rate. K252a (KI20), which consistently increased infection rates with both parasite forms, also increased schizont size in all isolates. A SI was calculated for size for each isolate in the same manner as was done for infection rates (**Fig. 6**). Using this metric, we again saw variation among isolates in sensitivity to kinase inhibition.

**Figure 6.**
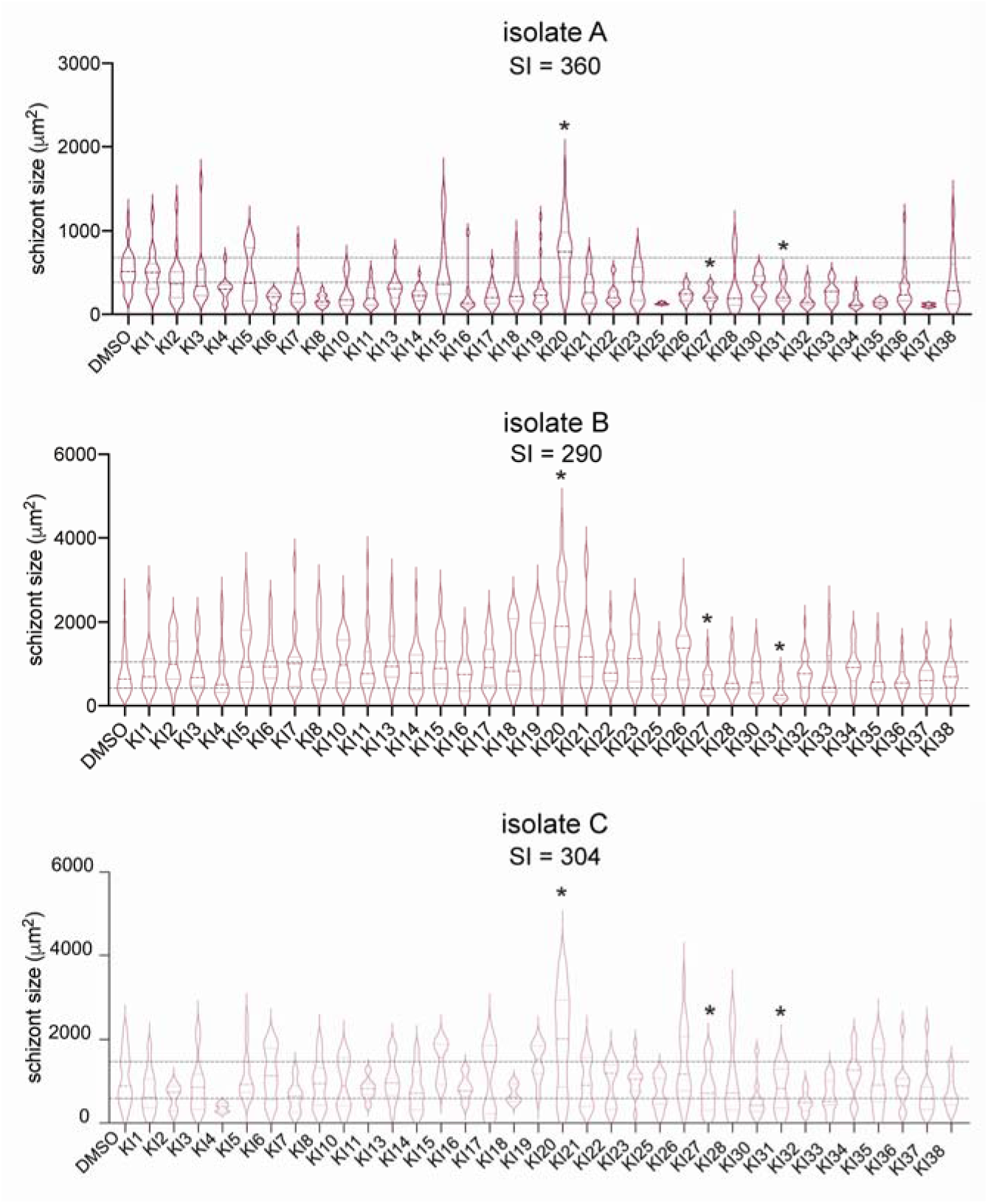
Host kinase inhibition influences schizont size. Effect of panel of 28 kinase inhibitors on schizont size for three different *P. vivax* isolates in primary human hepatocytes. Schizont area was quantified from 3 technical replicates for each inhibitor and normalized to a DMSO control 8 dpi. Inhibitors that consistently increased or decreased size across all 3 isolates, using a 20% cut-off, are highlighted. A sensitivity index (SI) was calculated for each isolate.

The effect of KIs on schizont size did correlate between each isolate pair, however this was strongly driven by the effect of K252a (KI20) (**Supplementary Fig. 5b**). A significant correlation existed between the effect of KIs on schizont infection and the effect on schizont size for isolates A and C, but not for isolate B (**Supplementary Fig, 5c**). These correlations were also heavily influenced by K252a (KI20) and only remained significant for isolate A when this inhibitor was removed. These data suggest that there may be distinct host factors that influence infection maintenance and parasite growth.

### Host kinase regulation of *P. vivax* LS infection across isolates and phenotypes

We next asked if the KI screen could identify host regulators of infection. We used KiR to predict host kinases whose activity impacts *P. vivax* schizont infection, hypnozoite infection, and schizont size. Data from each isolate (**Fig. 2, 6**) were run through KiR independently and kinases identified that were predicted for at least 2 of the 3 isolates were considered bona fide predictions (**Table 1, Supplementary Table 2**). One kinase, Erb-B2 receptor tyrosine kinase 2 (ERBB2), a member of the epidermal growth factor receptor family of receptor tyrosine kinases and an oncogene, was predicted to regulate all three phenotypes: schizont levels, hypnozoite levels, and schizont size across multiple isolates. Casein kinase 2 alpha 1 (CSNK2A1) was predicted to regulate both schizont and hypnozoite infection rates, however this could be confounded by a direct effect of inhibitors on the parasite casein kinase ^55,56^. Activin A receptor type 1B (ACVR1B), a growth and differentiation factor, and oxidative stress responsive kinase 1 (OXSR1), were predicted to regulate both schizont infection and size, consistent with previous work describing reactive oxygen species and lipid peroxidation regulation of *P. yoelii* infection and size ^29,57^. Two kinases were predicted to regulate both schizont size and hypnozoite infection, calcium/calmodulin-dependent protein kinase 4 (CAMK4) and dual specificity tyrosine phosphorylation regulated kinase 4 (DYRK4).

**Table 1.**
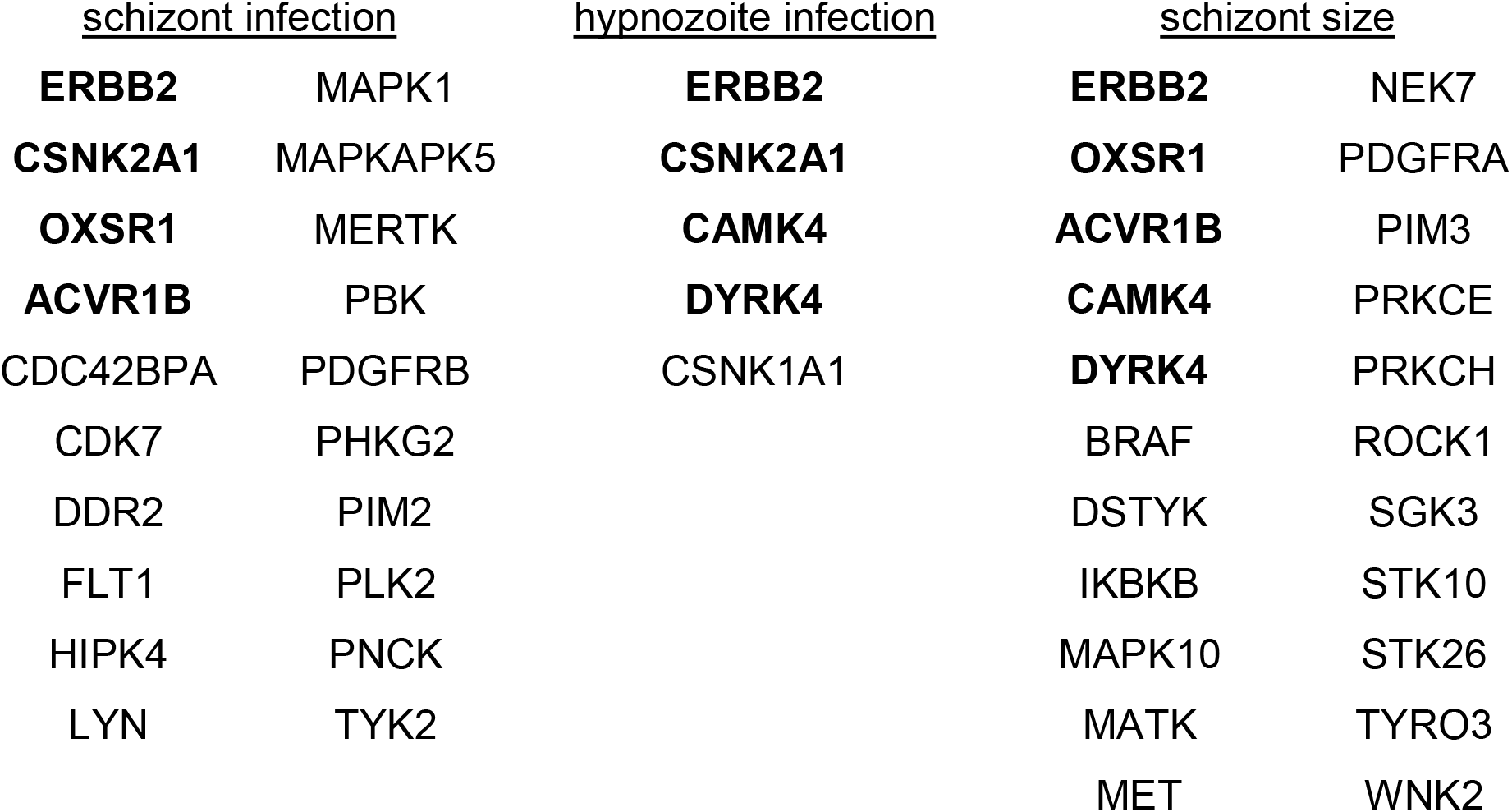
Host kinases predicted to regulate *P. vivax* infection. Host kinases predicted by KiR to regulate schizont infection, hypnozoite infection, or schizont size in at least two of three isolates. Overlapping kinases are in bold.

## Discussion

*Plasmodium vivax* is the malaria causing species with the greatest described diversity of phenotypic outcomes in the field. Even within sporozoites isolated from a single mosquito infection, some parasites rapidly develop in the liver and others remain non-dividing for weeks or months before eventual reactivation. The ratio and frequency of reactivation of these non-replicating parasites differs dramatically across geographic region and among isolates. Consistent with this observation, our data demonstrate a large amount of variation across *P. vivax* isolates in the hepatocyte phosphosignaling pathways that regulate their LS infection rates. Our data are consistent with the hypothesis that there is significant variation in the biology of *P. vivax* LS infection between individual parasite isolates collected from the same region, within the same CSP variant, and within the same lot of primary hepatocytes. Consistent with recently published work from Vantaux et al. ^24^, we found that parasite size, infection rate, and hypnozoite to schizont ratio varied significantly among parasite isolates under basal conditions, although we did observe more variation in schizont size among isolates within a single hepatocyte donor. Furthermore, we observed a large amount variation among *P. vivax* isolates in both the extent and specificity of their sensitivity to host kinase inhibition. Two parasite isolates that were processed in parallel did not show less variation compared to each other than compared to the isolate processed on a different date, suggesting that biological difference, rather than technical experimental variation, is driving the observed phenotypes. Significant heterogeneity has been described in genotype ^58,59^, transcriptional profiles ^39,60–63^, and infection phenotypes across the *P. vivax* lifecycle. Varying infection phenotypes include disease severity ^64,65^, selectivity index for reticulocytes ^66^, frequency of hypnozoite formation and timing of relapse ^5,11–13,17^. Our data suggest a potential molecular origin for some of these heterogeneous outcomes; that differing responses to host phosphosignaling could underly the infection phenotypes of different *P. vivax* isolates. Whether these differences in host phosphosignaling susceptibility extend beyond the liver stage of infection is an interesting area for future investigations.

Within the liver stage, the greater consistency (**Fig, 2,3**) and predictive power (**Fig. 5**) between schizont and hypnozoite forms within an isolate suggests that the genetic diversity between individual parasite isolates is a greater driver of phenotypic diversity in the response to host kinase inhibition than the differences between dormant and replicating parasite forms. Single cell transcriptional profiling of LS *P. vivax* and *P. cynomolgi* parasites identified multiple sub-clusters within schizont and hypnozoite populations ^39,63,67^. Subclusters of schizonts were posited to be defined by their pre-commitment, or lack thereof, to form gametocytes ^39^. Hypnozoite populations at a similar time point post-infection were also thought to form 3 clusters corresponding to persisting, early activated, and late activated states ^63^. Future work will be needed to determine the host-cell dependencies of these different parasite populations and to what extent parasite isolate influences their relative levels.

*Plasmodium* liver stage parasites are one of the fastest growing eukaryotes, yet very little is known about the intrinsic and environmental regulatory processes that control their size, although current data suggests that there are contributions from each. For example, host AMP-activated protein kinase (AMPK), which is likely inhibited by some drugs in our panel but is not part of the 300 kinases for which biochemical data is available ^41^, is linked to development of *P. berghei* and *P. falciparum* liver stage size. Manipulation of host AMPK activity had an inverse effect on LS size. Reduction in parasite size corresponded to less parasite replication and lead to reduced total merosome numbers, not simply a delay in development ^68^. Additionally, both host lipid uptake and regulation of host autophagic processes are required for optimal LS growth ^29,68,69^. There is also strong evidence for parasite-intrinsic regulators of growth. For example, inhibition of both host Tak1 and *P. falciparum* protein kinase 9 (PfPK9) increased the size of LS *P. berghei* ^70^. Additionally, parasite fatty acid synthesis pathways are required for complete growth and development in rodent infectious species of *Plasmodium* ^71^.

The interplay between host signaling and parasite size is of particular interest for *P. vivax,* due to its long residence in the liver and its ability to switch from a non-dividing to a rapidly developing form. One hypothesis is that growth is biologically linked to dormancy, and the molecular machinery that underlies hypnozoite formation also underlies schizont growth. This concept is not unprecedented in biology, as there are well-established linkages between cell size and cell division in other systems including bacteria ^72^, yeast ^73^, and several mammalian cell types ^74,75^. Consistent with this hypothesis, we found that the majority of the few host kinases that were predicted to regulate hypnozoite infection across isolates were also predicted to regulate schizont size (**Table 1**). Specifically, the overlapping kinases were ERBB2, an oncogene ^76^, CAMK4, which regulates transcription, protein synthesis, and apoptosis and has a role hepatocellular carcinoma growth ^77,78^, and DYRK4, which regulates differentiation, apoptosis and sensitivity to DNA damage ^79,80^. Investigating the link between dormancy and size, and how host regulators might influence parasite checkpoints in a way that is analogous to the established checkpoint model in other systems remains a critical area for future research.

While there is convincing evidence for parasite isolate determination of the ratio of schizont to hypnozoite formed upon infection ^5,11,12^, it remains an open question to what extent host conditions can skew this ratio. Our data support a role for host phosphosignaling in regulating the number of hypnozoites and schizonts present 8 dpi. We did not observe any kinase inhibitors that increased schizont while decreasing hypnozoite levels (**Fig. 3**), which would have supported a model of host regulated conversion between hypnozoite and schizont forms in the initial period of infection. Future work utilizing a time course of treatment and selective clearance of schizonts is warranted to further investigate the role of host phosphosignaling on hypnozoite formation and activation.

Genetic and epigenetic variation between and within patient isolates are likely candidates for the variation seen here. Future work incorporating more data on parasite genetic variation may confirm or refute the existence of consistent host kinase regulators of *P. vivax* infection. We anticipate that as new systems become available for studying *P. vivax* LS infection and more screens are conducted, comparisons and modeling between forms and isolates, as was done here, will add depth to our understanding of the complexities of infection and the interactions between host and parasite.

## Study Limitations

We attempted to minimize sources of variation by holding constant hepatocyte donor, CSP variant, province of isolate origin, and, for two of the three isolates, time of processing. However, we lack additional information about the source of the parasite isolates, particularly if they are from an acute or relapsing infection, that could explain isolate variation. The small number of parasite isolates available also limits our statistical power and ability to draw conclusions on the influence of isolate variation on susceptibility to host kinase inhibition.

Another limitation of this study is the inability to rule out off-target effects of inhibitors on parasite kinases. Inhibitors that influenced blood stage infection with *P. falciparum* were removed from the screen, however, different parasite kinases may be expressed in *P. vivax* liver-stage infection that cannot be accounted for. Future work using genetic approaches will be needed to validate host kinase involvement in any phenotype.

Additionally, the inhibitor screen used here only includes measured activity on 300 of the over 500 host kinases. When calculating SIs for each phenotype, the effects of inhibitors on kinases that are not part of the screen are not accounted for. As substrate activity screens of these inhibitors expand in the future, the additional data may be included in these indexes.

## Methods

### Generating *P. vivax* sporozoites

*Plasmodium vivax* infected blood was collected from patients attending malaria clinics in the Thasongyang district in Tak province, Thailand, under the protocol approved by the Ethics Committee of the Faculty of Tropical Medicine, Mahidol University (MUTM 2018-016-03). Written informed consent was obtained from each patient before sample collection. Infected blood was washed once with RPMI1640 incomplete medium before being resuspended with AB serum to a final 50% hematocrit and fed to female *Anopheles dirus* through membrane feeding. Engorged mosquitoes were maintained in 10% sugar solution until day 14-21 post feeding when salivary glands were dissected and sporozoites isolated. *P. vivax* mono-infection was confirmed by nested PCR. CSP variant was determined by RFLP-PCR.

### Infections

Primary human hepatocytes (lot ZPE, BioIVT) were plated in collagen-coated 384-well plates at 25,000 cells per well as described previously ^50^. A 3-well border of water was used around each plate to minimize edge effects. Two days post-plating, cells were infected with 14,000 *P. vivax* sporozoites per well. Wells were treated with an inhibitor or DMSO control 24h post-infection. Media was changed and treatments refreshed every other day until fixation in 4% PFA on day 8 post-infection. Cells were permeabilized and blocked with 1% TritonX-100 and 2% BSA, and then stained with DAPI and an antibody against PvUIS4 ^53^.

### Imaging and quantification

Plates were imaged using a Keyence BZ-X700 and quantified using ImageJ. Parasites were identified by positive PvUIS4 and DAPI staining and were categorized as schizonts (diameter > 10μm) or hypnozoites (diameter < 10μm, UIS4 prominence). To evaluate edge effects parasite levels were compared between outer (D5-15, L5-15) and inner (F5-15, J5-15) wells that received the same treatment. Schizont area was measured using PvUIS4 staining with ImageJ. Hepatocyte nuclei were counted using ImageJ as follows: DAPI channel image threshold was set with a pixel value lower limit of 69 and upper limit of 255. Nuclei were quantified using the measure particles feature within a size range of 40-800 pixel units and circularity between 0.4-1 to exclude debris and schizont DNA.

### Sensitivity index

To create a sensitivity index (SI) for each phenotype a series of independent, random weights ranging from 0 and 1 were generated for the 34 kinase inhibitors in the screen. For each of the 300 kinases in the dataset, the sum of the weighted residual activity (catalytic activity upon kinase inhibitor treatment) from all 34 kinases inhibitors was calculated, after which, standard deviation of the summed residual activity across the 300 kinases was calculated. The above procedure was repeated for 100,000 iterations, and the weights that led to the minimum standard deviation were assigned to the kinase inhibitors. The effect of each kinase inhibitor (percent change compared to DMSO) was then multiplied by the assigned weight. These weighted values were then added together to create the SI for each phenotype.

### Kinase Regression predictions

Kinase regression was performed independently for each parasite isolate and each phenotype, using the previously published algorithm described by Arang et al. ^48^. Glmnet for Python (version 2.2.1) was used to fit the generalized linear models via penalized maximum likelihood, with the elastic net mixing parameter α of 0.8. The kinases with non-zero coefficients were predicted to be important for the phenotype of interest, and the effect of kinase inhibitors not tested in this study on that phenotype was also predicted based on the kinase predictions.

### Statistics

Distributions of all data sets were analyzed for normality using the Anderson-Darling and D’Agostino & Pearson tests. Infection rate and form ratio data were normally distributed and analyzed by ANOVA with Tukey’s multiple comparisons test. Schizont size distributions were not normally distributed among all isolates and were analyzed using the Kruskal-Wallis test with Dunn’s multiple comparisons test. To compare variances in schizont size between isolates, the data were ln(X) transformed before being analyzed by F test. Toxicity data were analyzed for all replicates by two-way ANOVA and Dunnett’s multiple comparisons test. Paired t-tests were used to test for edge effects on infection. Inhibitor dose response data were analyzed by Fisher’s LSD test. Pearson correlation coefficients were used to analyze all correlation data.

## Supporting information

Supplementary File 1 - Raw data

## Acknowledgements

This work was funded by R21AI151344 and R01AI177257 from the National Institutes of Health to AK and EKKG.

## Author Contributions

Conceptualization, AK and EKKG; Methodology, AK and EKKG; Investigation, EKKG, LW, VIP, TT, and CY. Resources, WR and JS. Writing – Original Draft, EKKG. Writing – Reviewing and Editing, EKKG, AK, LW, VIP, TT, CY, WR, and JS; Funding Acquisition, AK, EKKG, and JS; Supervision, AK and JS.

## Supplemental Material

**Supplementary Figure 1.**
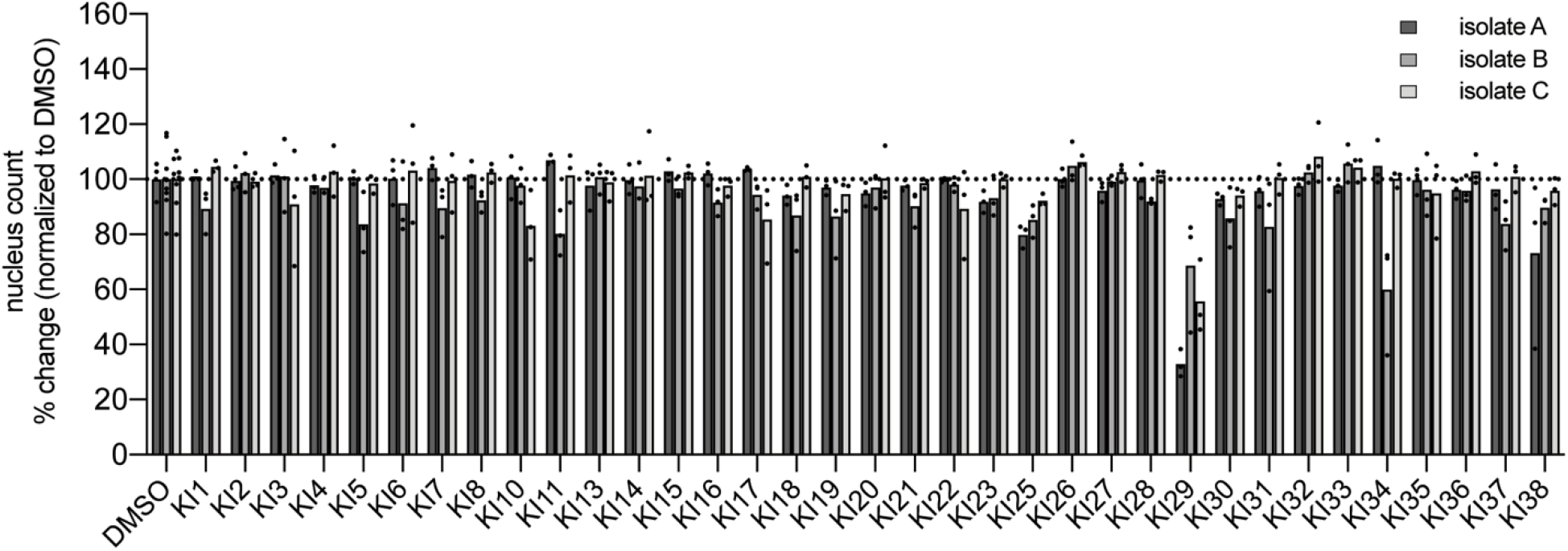
Toxicity of kinase inhibitors in primary hepatocytes. Primary hepatocyte nuclei were quantified in each infected well 8 dpi at the time of parasite quantification as a measure of kinase inhibitor toxicity. Each dot represents a technical replicate. N = 3-9. Data were normalized to the DMSO control average for each isolate and analyzed by Dunnett multiple comparisons test.

**Supplementary Figure 2.**
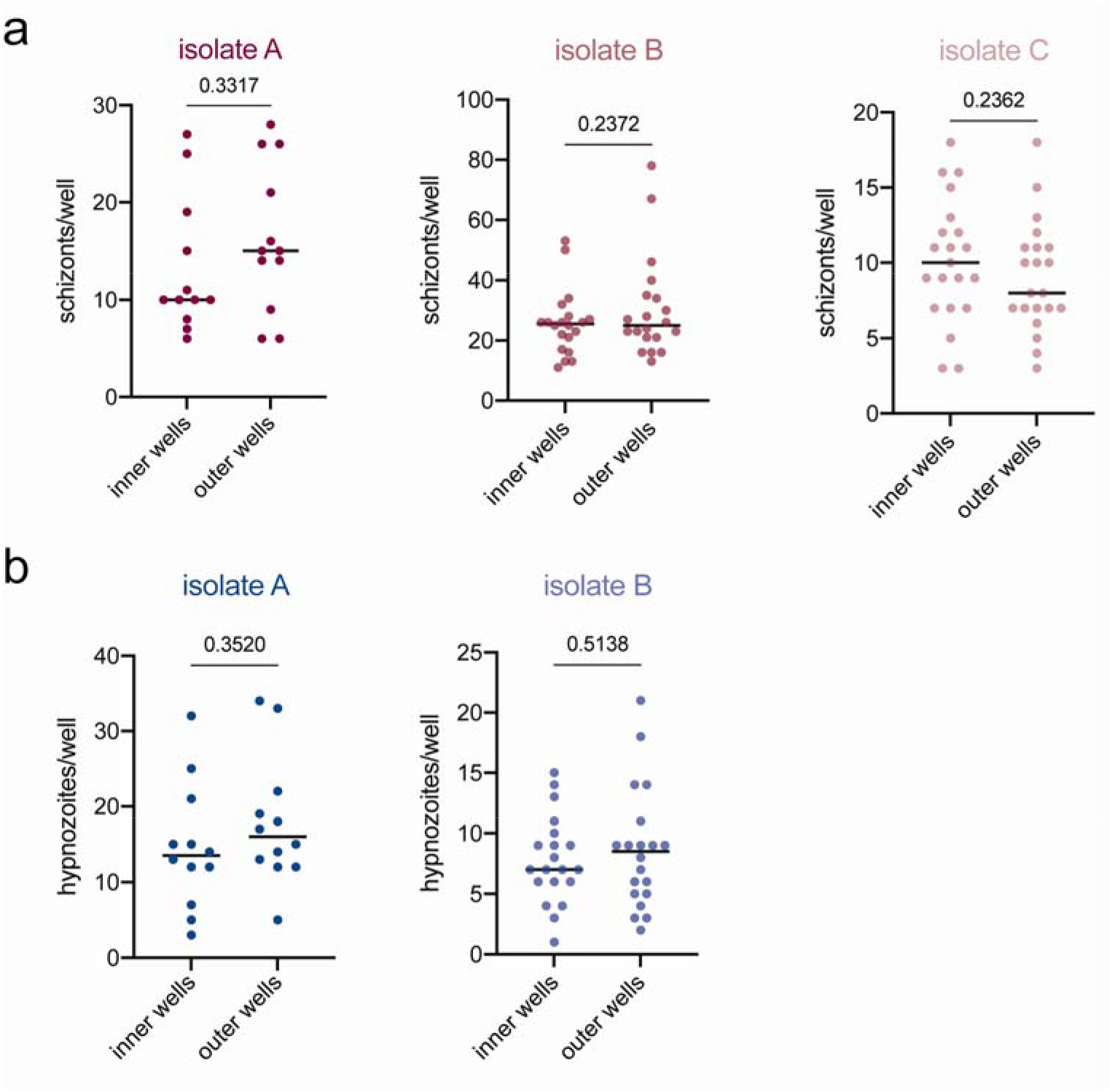
Well position did not influence infection rates. (A) Schizont and (B) hypnozoite infection rates were compared between outer edge (D5-15, L5-15) and inner (F5-15, J5-15) wells in each 384-well plate (one plate per isolate) that received the same treatment. Each dot represents a well. Data were analyzed by paired t-test.

**Supplementary Figure 3.**
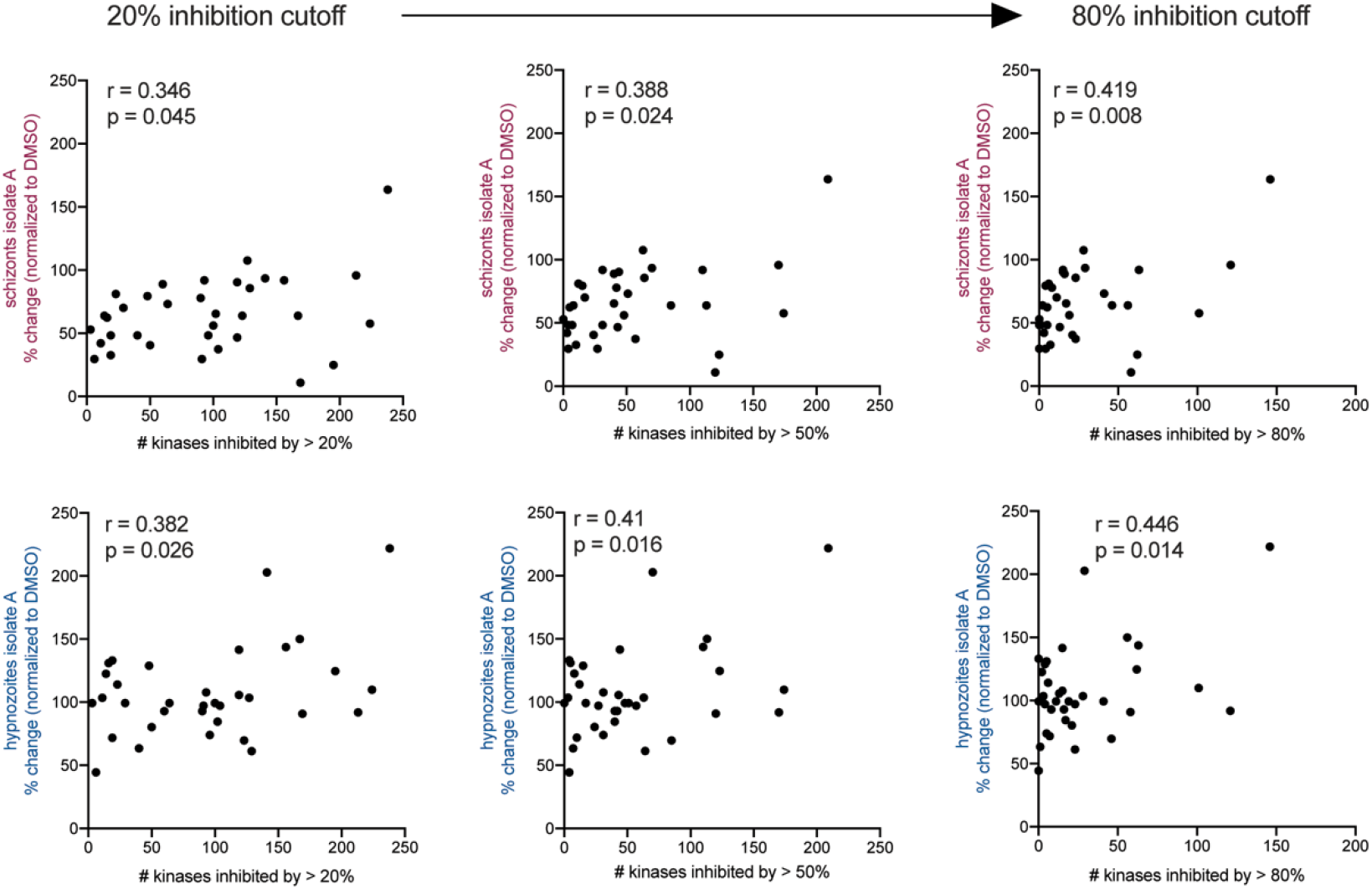
Promiscuous inhibitors do not show greater inhibition of infection. The effect of each inhibitor on schizont or hypnozoite infection, for isolate A, is plotted against the number of kinases inhibited by each KI 41 using cut-offs of 20%, 50% or 80%. Each dot represents a single inhibitor. Pearson correlation coefficients (r) and p-values were calculated for each plot.

**Supplementary Figure 4.**
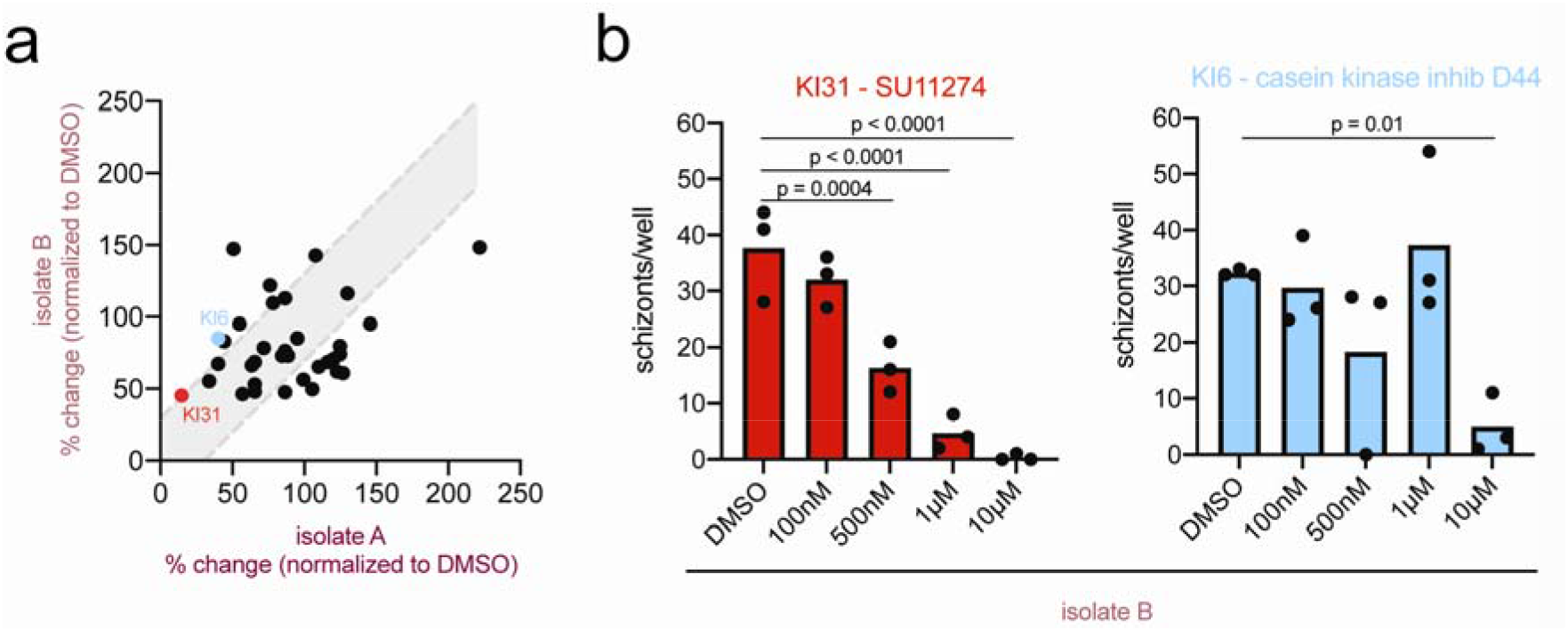
Schizont sensitivity to inhibitors varies across isolates. (A) Correlation between effect of kinase inhibitors on schizont infection compared to controls in isolate A vs isolate B. Each dot represents an inhibitor. (B) Schizont infection dose response curves for two kinase inhibitors. Each dot represents a technical replicate. Data were analyzed by Fisher’s LSD test.

**Supplementary Figure 5.**
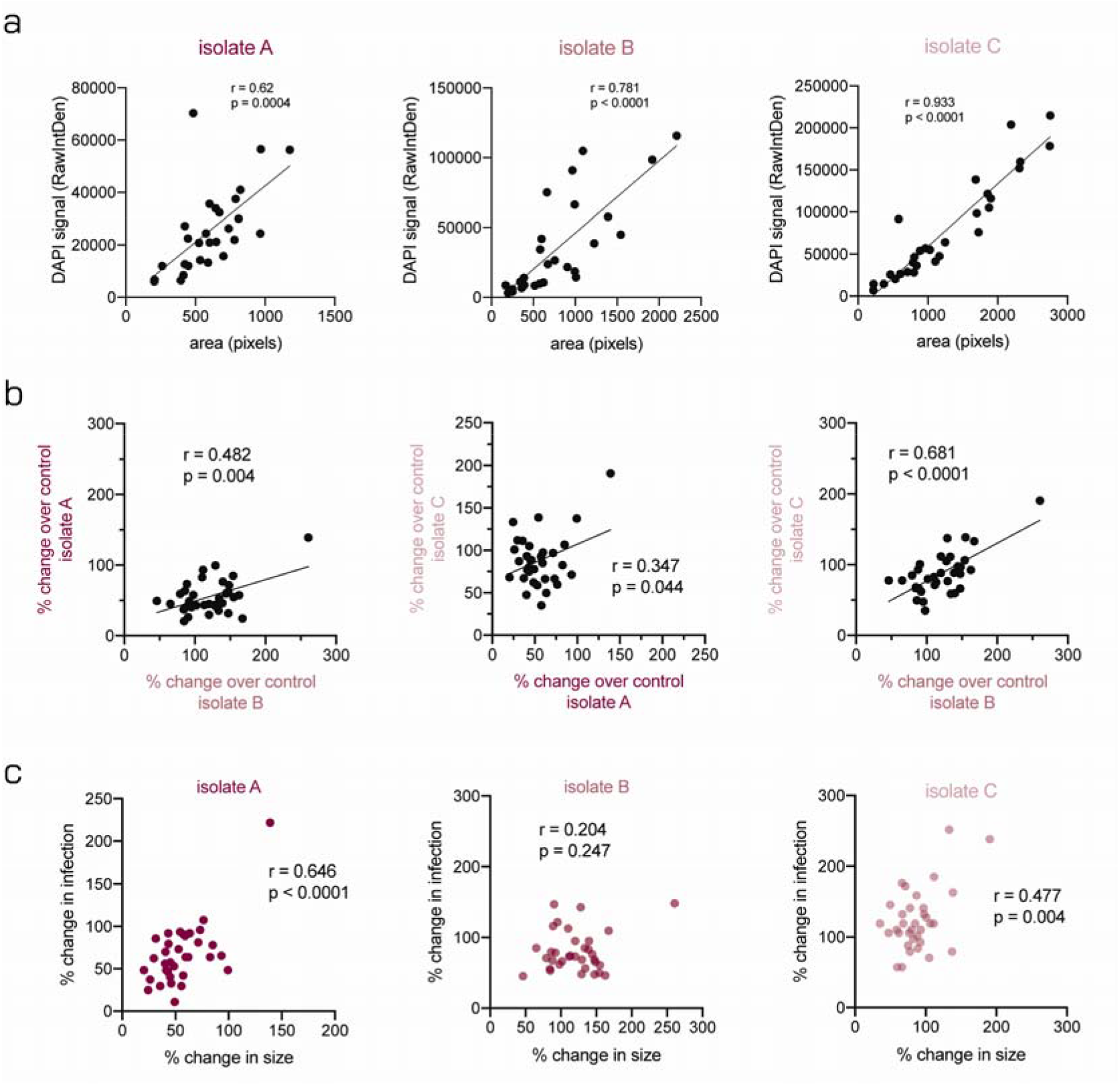
Correlation of effect on size between isolates and with infection. (A) Schizont size plotted against raw density of DAPI signal within the parasite in control wells. Each dot represents a single parasite from a DMSO-treated well. (B) The effect of kinase inhibitors on parasite size plotted for each pair of isolates. Each dot represents a kinase inhibitor. (C) Effect of kinase inhibitors on parasite size plotted against effect on schizont infection rate for each isolate. Each dot represents an inhibitor. Pearson correlation coefficients (r) and p-values (p) were calculated for each comparison.

