## Supplementary Table 1 for "Host kinase regulation of *Plasmodium vivax* dormant and replicating liver stages"

| KI # | Name | CAS | weight |
|---|---|---|---|
| 1 | Aminopurvanolol A | 220792-57-4 | 0.51 |
| 2 | GSK inhibitor IX (BIO) | 667463-62-9 | 0.2 |
| 3 | Bosutinib | 380843-75-4 | 0.05 |
| 4 | CDK2 inhibitor IV; NU6140 | 444723-13-1 | 0.45 |
| 5 | CDK4 inhibitor | 546102-60-7 | 0.98 |
| 6 | Casein kinase I inhibitor D44 | 301836-43-1 | 0.79 |
| 7 | Dasatinib | 183319-69-9 | 0.42 |
| 8 | AMPK Inhibitor; Compound C (Dorsomorphin) | 866405-64-3 | 0.04 |
| 9 | Dovitinib | 405169-15-6 | *omitted due to blood sta |
| 10 | EGFR/ErbB2/ErbB4 inhibitor | 881001-19-0 | 0.52 |
| 11 | Erlotinib | 184475-35-2 | 0.05 |
| 12 | Gefitinib | 220127-57-1 | *omitted due to blood sta |
| 13 | Go 6976 | 136194-77-9 | 0.15 |
| 14 | Go 6983 | 133053-19-7 | 0.4 |
| 15 | GSK-3 Inhibitor X | 740841-15-0 | 0.09 |
| 16 | GSK-3 Inhibitor XIII | 404828-08-6 | 0.05 |
| 17 | H89 | 127243-85-0 | 0.26 |
| 18 | Imatinib | 641571-10-0 | 0.18 |
| 19 | JAK inhibitor I | 457081-03-7 | 0.17 |
| 20 | K252a | 97161-97-2 | 0.04 |
| 21 | Lapatinib | 231277-92-2 | 0.39 |
| 22 | Lck inhibitor | 213743-31-8 | 0.17 |
| 23 | Masitinib | 790299-79-5 | 0.27 |
| 24 | Nilotinib | 443913-73-3 | *omitted due to blood sta |
| 25 | PKR inhibitor | 608512-97-6 | 0.18 |
| 26 | SB218078 | 135897-06-2 | 0.24 |
| 27 | Sorafenib | 284461-73-0 | 0.62 |
| 28 | JNK inhibitor II (SP600125) | 129-56-6 | 0.22 |
| 29 | Staurosporine | 62996-74-1 | omitted due to hepatocyt |
| 30 | Staurosporine n benzoyl | 120685-11-2 | 0.4 |
| 31 | SU11274 | 658084-23-2 | 0.01 |
| 32 | SU6656 | 330161-87-0 | 0.44 |
| 33 | GSK-3b inhibitor I (TDZD-6) | 327036-89-5 | 0.74 |
| 34 | Tofacitnib | 477600-75-2 | 0.46 |
| 35 | TWS119 | 601514-19-6 | 0.02 |
| 36 | Vandetanib | 146986-50-7 | 0.42 |
| 37 | ROCK inhibitor (Y-27632) | 146986-50-7 | 0 |
| 38 | Cdk1/2 Inhibitor III | 443798-55-8 | 0.01 |

ge reactivity

ge reactivity

ge reactivity
