## Supplementary Table 2 for "Host kinase regulation of *Plasmodium vivax* dormant and replicating liver stages"

**Isolate A**

**kinase predictions**

**Schizonts**

| Gene ID | Coefficient |
|---|---|
| ACVR1B | 0.17363098 |
| CDC42BPA | 0.21804341 |
| CLK4 | 0.17296438 |
| CSNK1A1 | -0.0326162 |
| CSNK2A1 | 0.18820304 |
| DDR2 | -0.0016315 |
| EPHA8 | 0.08181448 |
| ERBB2 | 0.02122199 |
| FGFR2 | -0.2482772 |
| FLT1 | -0.0593492 |
| GRK4 | -0.4755787 |
| GRK5 | 0.48142834 |
| HIPK4 | -0.1285024 |
| LYN | -0.0294927 |
| MAP2K1 | -0.3222146 |
| MAP3K14 | -0.3637078 |
| MAPK12 | 1.12061785 |
| MAPKAPK5 | 0.36480352 |
| MATK | 0.23606743 |
| NEK3 | -0.1883369 |
| NLK | 0.35619389 |
| OXSR1 | -0.2821611 |
| PAK6 | -0.0478354 |
| PBK | 0.01349312 |
| PDGFRB | 0.00190909 |
| PHKG2 | -0.0002235 |
| PIM2 | 0.0652672 |
| PRKACA | -0.3996053 |
| PRKCE | 0.0604892 |
| PTK2 | -0.167157 |
| PTK6 | -0.1116583 |
| RIPK2 | -0.1435123 |
| STK10 | -0.0724907 |
| TSSK1B | 0.13606736 |
| TSSK2 | -0.0112251 |
| TYK2 | -0.0806468 |

**hypnozoites**

| Gene ID | Coefficient |
|---|---|
| AKT1 | -0.5497513 |
| BRAF | -0.3419016 |
| CAMK4 | 0.87451717 |
| CHUK | 0.20006266 |
| CSK | -0.0043528 |
| CSNK1A1 | -0.1932317 |
| CSNK2A1 | 0.36171118 |
| DMPK | 0.60594185 |
| DYRK4 | 0.58763392 |
| EPHB4 | 0.16590199 |
| ERBB2 | 0.00804346 |
| ERBB4 | 0.13491547 |
| FLT1 | -0.4574981 |
| GRK3 | 0.5522592 |
| GRK5 | 0.2064367 |
| GSK3A | 0.1129771 |
| HIPK4 | -0.6863241 |
| JAK1 | -0.0669944 |
| MAP3K14 | -0.3262994 |
| MST1R | 0.13134244 |
| NEK11 | -0.0566966 |
| NLK | 0.07596804 |
| OXSR1 | 0.38301203 |
| PAK2 | -0.1146037 |
| PIM3 | 0.12466191 |
| PRKACA | -0.2137447 |
| PRKCZ | -0.0619058 |
| PTK2B | -0.1497714 |
| SRPK2 | -0.2975894 |
| STK11 | -0.0806712 |
| STK25 | 0.2032991 |
| TAOK2 | -0.0562455 |
| TBK1 | 0.01277198 |
| TEK | -0.0183146 |
| TYK2 | -0.0095873 |
| TYRO3 | -0.2758903 |
| WNK2 | 0.49379627 |

**Schizonts**

| Drug |
|---|
| PKR_Inhb_Neg_Cntr |
| VX-702 |
| VEGFRTK_Inhb_III_KRN633 |
| CaseinK_II_Inhb_III_TBCA |
| Nilotinib |
| DNA-PK_Inhb_III |
| PI_3-Kg_Inhb |
| PP3 |
| PI_3-Kg_Inhb_II |
| JNK_Inhb_V |
| AG_1295 |
| p38_MAPK_Inhb |
| Roscovitine |
| TGF-b_RI_Inhb_III |
| MetK_Inhb |
| PD_169316 |
| JNK_Inhb_IX |
| Cdk2_Inhb_III |
| AG_9 |
| Imatinib |
| JAK3_Inhb_IV |
| DNA-PK_Inhb_II |
| p38_MAPK_Inhb_III |
| Lapatinib |
| Cdk4_Inhb |
| MNK1_Inhb |
| LY_303511-_Negative_control |
| PKCbII/EGFR_Inhb |
| JNK_Inhb_VIII |
| GSK3_Inhb_X |
| Tandutinib |
| Alsterpaullone_2Cyanoethyl |
| BAY_11_7082 |
| JNK_Inhb_Neg_Cntr |
| EGFR_Inhb |
| MEK_Inhb_I |
| PDGFRTK_Inhb_III |
| AG_490 |
| IC261 |

PI-103
Bisindolylmaleimide_I
PDGFRTK_Inhb_IV
SB220025
PKR_Inhb
AG_1296
ROCK_InhbY
LY294002
MEK1/2_Inhb
Herbimycin_A_Streptomyces_sp.
Compound_52
VEGFR2K_Inhb_IV
Flt-3_Inhb
Dasatinib
Fascaplysin_Synthetic
NF-kB_Activation_Inhb
VEGFR2K_Inhb_II
Syk_Inhb_II
Cdk1/5_Inhb
Wortmannin
Mubritinib
Vatalanib
GSK3b_Inhb_XI
JAK_Inhb_I
Cdk2_Inhb_IV_NU6140
SU9516
SKF-86002
GSK3b_Inhb_II
STO-609
Ro-32-0432
Flt-3_Inhb_II
VEGFRTK_Inhb_II
Masitinib
Kenpaullone
PP1_Analog_II
PKCb_Inhb
Aurora/Cdk_Inhb
PD_158780
PDGFRTK_Inhb_II
Akt_Inhb_VIII_Isozyme-Selective_
Isogranulatimide
Chelerythrine_Chloride
KN-62

MK2a_Inhb
Tpl2_K_Inhb
Cdk1_Inhb_CGP74514A
SB202474
IGF-1R_Inhb_II
TGF-b_RI_K_Inhb
MEK_Inhb_II
GSK3_Inhb_XIII
IRAK-1/4_Inhb
Erlotinib
Chk2_Inhb_II
KN-93
PDK1/Akt/Flt
JNK_Inhb_II
ERK_Inhb_III
Gefitinib
AMPK_Inhb_Comp_C
Diacylglycerol_K_Inhb_II
DNA-PK_Inhb_V
cFMS_RTK_Inhb
AG_112
Flt-3_Inhb_III
Syk_Inhb_III
SB203580
Vandetanib
Tofacitinib
Akt_Inhb_X
Rho_K_Inhb_III_Rockout
Purvalanol_A
Cdk4_Inhb_II_NSC_625987
Akt_Inhb_V_Triciribine
PD_98059
JAK3_Inhb_VI
Indirubin_E804
Aminopurvalanol_A
Rapamycin
PD_174265
SB202190
G_6983
Sphingosine_K_Inhb
DMBI
Cdc2-LikeK_Inhb_TG003
Bohemine

ERK_Inhb_II_FR180204
SC-68376
G_6976
Tozasertib
AGL_2043
Bisindolylmaleimide_IV
VEGF_Receptor_2_K_Inhb_I
Indirubin-3′-monoxime
Compound_56
JAK3_Inhb_II
HA_1077_Dihydrochloride_Fasud
GSK3b_Inhb_I
Cdk/Crk_Inhb
Rho_K_Inhb_IV
GSK3b_Inhb_VIII
Akt_Inhb_IV
GTP-14564
AG_1478
Staurosporine_N
PDGFRTK_Inhb
Aloisine_RP106
EGFR/ErbB-2_Inhb
EGFR/ErbB-2/4_Inhb
Bcr-abl_Inhb
Sunitinib
Aurora_Inhb_III
Alsterpaullone
Cdk1_Inhb
Pazopanib
Sorafenib
Cdk4_Inhb_III
CaseinK_I_Inhb_D44
BPIQ-I
AG_1024
ATM/ATR_K_Inhb
VEGFR2K_Inhb_III
ATM_K_Inhb
Syk_Inhb
GSK3_Inhb_IX
LCK_Inhb
Aloisine_A_RP107
Dovitinib
IKK-2_Inhb_IV

SrcK_Inhb_I
SU11652
SB218078
Bosutinib
Cdk1/2_Inhb_III
SU6656
Staurosporine
H-89_Di
TWS119
K-252a

**drug predictions**

| Prediction | Drug | Prediction |
|---|---|---|
| | **Hypnozoites** | |
| 21.9336765 | JNK_Inhb_V | 0.06309739 |
| 29.549652 | Staurosporine | 8.49962448 |
| 30.4512892 | Alsterpaullone_2Cyanoethyl | 13.4944128 |
| 35.5886945 | Alsterpaullone | 19.1180133 |
| 37.741286 | Bisindolylmaleimide_I | 24.7024328 |
| 38.9900064 | Aloisine_RP106 | 25.9776593 |
| 39.1995853 | Bcr-abl_Inhb | 30.8224095 |
| 40.0851166 | AG_1295 | 31.5908324 |
| 42.6328225 | PI_3-Kg_Inhb_II | 32.1415102 |
| 42.9857576 | CaseinK_II_Inhb_III_TBCA | 34.2204541 |
| 44.473398 | Tpl2_K_Inhb | 37.6306396 |
| 45.2219666 | Cdk1/5_Inhb | 39.2480951 |
| 46.3109936 | Cdk4_Inhb | 40.1166801 |
| 46.6935821 | Dasatinib | 40.8491277 |
| 47.5949444 | TGF-b_RI_K_Inhb | 41.7767675 |
| 47.6565065 | Erlotinib | 42.0171284 |
| 47.8248759 | Gefitinib | 43.5088272 |
| 48.0103273 | Bisindolylmaleimide_IV | 44.5194687 |
| 48.9436702 | GSK3_Inhb_XIII | 44.7178411 |
| 48.9672336 | Aloisine_A_RP107 | 44.84552 |
| 49.3123182 | Akt_Inhb_X | 45.1166186 |
| 49.3453327 | ATM_K_Inhb | 45.8273083 |
| 49.4795148 | AG_490 | 46.0395709 |
| 49.4839212 | GSK3_Inhb_X | 46.4935455 |
| 49.4852013 | BAY_11_7082 | 46.6529186 |
| 49.6823993 | JAK3_Inhb_IV | 47.7205023 |
| 49.7717494 | ROCK_InhbY | 48.0507303 |
| 49.7856537 | ATM/ATR_K_Inhb | 48.3627591 |
| 50.2617511 | Roscovitine | 48.5488074 |
| 50.3752363 | Compound_56 | 48.7000839 |
| 50.7860339 | Aurora_Inhb_III | 49.5396353 |
| 50.9229336 | GSK3b_Inhb_XI | 50.3151276 |
| 51.2053189 | DNA-PK_Inhb_II | 50.5011752 |
| 51.3071316 | Akt_Inhb_IV | 50.6298318 |
| 53.2010537 | AMPK_Inhb_Comp_C | 51.2631313 |
| 53.4135036 | Rho_K_Inhb_IV | 51.4730317 |
| 53.8191396 | Cdk1_Inhb_CGP74514A | 52.1997141 |
| 54.2548699 | PKCb_Inhb | 52.3579363 |
| 54.8043369 | Tandutinib | 52.6448527 |

**kinas**

**Schizonts**

| Gene ID | Coefficient |
|---|---|
| ACVR1B | 0.17363098 |
| CDC42BPA | 0.21804341 |
| CLK4 | 0.17296438 |
| CSNK1A1 | -0.0326162 |
| CSNK2A1 | 0.18820304 |
| DDR2 | -0.0016315 |
| EPHA8 | 0.08181448 |
| ERBB2 | 0.02122199 |
| FGFR2 | -0.2482772 |
| FLT1 | -0.0593492 |
| GRK4 | -0.4755787 |
| GRK5 | 0.48142834 |
| HIPK4 | -0.1285024 |
| LYN.1 | -0.0294927 |
| MAP2K1 | -0.3222146 |
| MAP3K14 | -0.3637078 |
| MAPK12 | 1.12061785 |
| MAPKAPK5 | 0.36480352 |
| MATK | 0.23606743 |
| NEK3 | -0.1883369 |
| NLK | 0.35619389 |
| OXSR1 | -0.2821611 |
| PAK6 | -0.0478354 |
| PBK | 0.01349312 |
| PDGFRB | 0.00190909 |
| PHKG2 | -0.0002235 |
| PIM2 | 0.0652672 |
| PRKACA | -0.3996053 |
| PRKCE | 0.0604892 |
| PTK2 | -0.167157 |
| PTK6 | -0.1116583 |
| RIPK2 | -0.1435123 |
| STK10 | -0.0724907 |
| TSSK1B | 0.13606736 |
| TSSK2 | -0.0112251 |
| TYK2 | -0.0806468 |

| | | |
|---|---|---|
| 55.0800941 | PKR_Inhb | 52.6686287 |
| 55.1510192 | JNK_Inhb_VIII | 53.1149832 |
| 55.1692983 | Cdc2-LikeK_Inhb_TG003 | 53.5009139 |
| 55.273327 | STO-609 | 53.6326193 |
| 55.3469557 | VX-702 | 53.6753273 |
| 55.4228823 | PI_3-Kg_Inhb | 53.6756323 |
| 55.4492374 | NF-kB_Activation_Inhb | 55.3356951 |
| 56.085314 | Akt_Inhb_VIII_Isozyme-Selecti | 55.3750906 |
| 56.1318179 | LY_303511-_Negative_control | 55.6052176 |
| 56.1863597 | Cdk2_Inhb_IV_NU6140 | 56.2601119 |
| 56.3088307 | p38_MAPK_Inhb | 56.6065027 |
| 57.3135164 | GSK3b_Inhb_I | 57.3130805 |
| 57.9157648 | p38_MAPK_Inhb_III | 57.3515645 |
| 58.3570047 | Cdk/Crk_Inhb | 57.5672723 |
| 58.3595588 | Ro-32-0432 | 57.6287981 |
| 58.4251262 | Bohemine | 57.7131497 |
| 59.3818613 | Lapatinib | 57.7690389 |
| 59.4374586 | AG_1024 | 57.8919277 |
| 59.5458945 | Flt-3_Inhb | 58.1914854 |
| 59.9277953 | Cdk1_Inhb | 58.9286296 |
| 60.4485727 | VEGFR2K_Inhb_II | 59.065591 |
| 60.4801253 | Sphingosine_K_Inhb | 59.3304089 |
| 60.5547176 | Mubritinib | 59.4227344 |
| 61.459063 | Cdk4_Inhb_II_NSC_625987 | 59.5304789 |
| 61.9603875 | AG_112 | 60.3122065 |
| 62.1122355 | Kenpaullone | 60.4278262 |
| 62.1479376 | AGL_2043 | 60.6413845 |
| 62.2935962 | Isogranulatimide | 60.7937243 |
| 62.6592338 | Chelerythrine_Chloride | 61.0252852 |
| 62.6965073 | DNA-PK_Inhb_V | 61.3883119 |
| 62.8064962 | JNK_Inhb_Neg_Cntr | 61.4344367 |
| 63.8796487 | TGF-b_RI_Inhb_III | 61.54287 |
| 64.1543247 | Chk2_Inhb_II | 61.6623911 |
| 64.3418854 | GSK3b_Inhb_VIII | 61.9759168 |
| 64.4845927 | DNA-PK_Inhb_III | 62.2092207 |
| 64.5422853 | GSK3b_Inhb_II | 62.7361537 |
| 64.9272206 | Cdk4_Inhb_III | 62.8640495 |
| 64.9447432 | CaseinK_I_Inhb_D44 | 62.9772508 |
| 65.0190744 | LY294002 | 63.1500396 |
| 65.160165 | Masitinib | 63.3351803 |
| 65.3760286 | Syk_Inhb_III | 63.5803401 |
| 65.8830586 | ERK_Inhb_II_FR180204 | 63.8461473 |
| 65.9982657 | KN-93 | 63.9344906 |

| | | |
|---|---|---|
| 66.0390717 | Wortmannin | 64.0726025 |
| 66.164004 | SU9516 | 64.3351865 |
| 66.385087 | EGFR/ErbB-2_Inhb | 64.4886396 |
| 66.6512513 | ERK_Inhb_III | 64.7054802 |
| 66.7329267 | Akt_Inhb_V_Triciribine | 64.8019045 |
| 67.0467513 | Imatinib | 65.0125395 |
| 67.1519884 | JNK_Inhb_II | 66.2186294 |
| 67.4479099 | Indirubin_E804 | 66.3564885 |
| 67.562018 | EGFR/ErbB-2/4_Inhb | 66.8642041 |
| 67.6624445 | VEGF_Receptor_2_K_Inhb_I | 67.1020951 |
| 67.6791053 | Tozasertib | 67.1822257 |
| 67.7260872 | Rho_K_Inhb_III_Rockout | 67.9707921 |
| 67.8543171 | PKR_Inhb_Neg_Cntr | 68.0918683 |
| 67.919288 | MEK_Inhb_II | 68.2828156 |
| 68.5565862 | SrcK_Inhb_I | 68.7139422 |
| 68.7713271 | JAK_Inhb_I | 69.1341107 |
| 68.8317316 | Syk_Inhb_II | 69.5378602 |
| 69.3399684 | SB202474 | 69.5604833 |
| 69.5397076 | IGF-1R_Inhb_II | 70.0915612 |
| 69.8949499 | DMBI | 70.092429 |
| 69.909924 | BPIQ-I | 70.0985554 |
| 70.022441 | Syk_Inhb | 70.1198943 |
| 70.1777999 | HA_1077_Dihydrochloride_Fa | 70.1758344 |
| 70.240691 | Herbimycin_A_Streptomyces_ | 70.4609487 |
| 70.8048491 | PD_158780 | 70.6193677 |
| 71.3865386 | Purvalanol_A | 71.0628471 |
| 71.4448952 | Compound_52 | 71.2070597 |
| 71.4681118 | SB203580 | 71.4241107 |
| 71.6737273 | Vatalanib | 71.4249744 |
| 71.686612 | cFMS_RTK_Inhb | 72.0403715 |
| 72.2355575 | G_6976 | 72.0839909 |
| 72.3796906 | JAK3_Inhb_VI | 72.1020885 |
| 72.6956495 | Cdk2_Inhb_III | 72.2749201 |
| 73.0778303 | Rapamycin | 72.3480595 |
| 73.2491118 | AG_9 | 72.6353418 |
| 73.3673178 | JNK_Inhb_IX | 72.674416 |
| 73.7239487 | Nilotinib | 72.8087342 |
| 74.130054 | Vandetanib | 72.859401 |
| 74.1635468 | G_6983 | 72.9501463 |
| 74.3221134 | IC261 | 73.5865134 |
| 74.8920993 | Flt-3_Inhb_II | 73.8728023 |
| 74.9052657 | PDGFRTK_Inhb_II | 74.7052218 |
| 75.0418458 | LCK_Inhb | 74.7524262 |

| | | |
|---|---|---|
| 75.250361 | JAK3_Inhb_II | 74.7756938 |
| 75.8798092 | Diacylglycerol_K_Inhb_II | 74.7773458 |
| 75.9927555 | MEK1/2_Inhb | 75.4767097 |
| 76.0514734 | KN-62 | 75.6636006 |
| 76.3609318 | Tofacitinib | 76.3492307 |
| 76.7814489 | SC-68376 | 76.3600781 |
| 77.2770423 | Indirubin-3′-monoxime | 76.8476684 |
| 77.2936369 | AG_1296 | 77.0242988 |
| 77.3426094 | SB220025 | 77.4903048 |
| 77.4824041 | PP3 | 78.6631485 |
| 77.6678553 | PI-103 | 78.8427986 |
| 78.0610082 | MEK_Inhb_I | 78.9149695 |
| 78.7463202 | PP1_Analog_II | 79.3508386 |
| 79.4689437 | PDGFRTK_Inhb | 79.7385078 |
| 79.9580993 | SKF-86002 | 80.2140604 |
| 80.0385871 | VEGFR2K_Inhb_III | 81.5469691 |
| 80.5563566 | Aminopurvalanol_A | 81.6409521 |
| 80.6193964 | AG_1478 | 81.6625421 |
| 80.8065188 | EGFR_Inhb | 81.8942131 |
| 81.7992824 | VEGFRTK_Inhb_III_KRN633 | 82.0428312 |
| 82.2770518 | IKK-2_Inhb_IV | 82.7825729 |
| 82.4107196 | PD_174265 | 83.2994283 |
| 82.4594019 | IRAK-1/4_Inhb | 84.133721 |
| 82.9243244 | MK2a_Inhb | 84.5438316 |
| 83.0714369 | PDGFRTK_Inhb_III | 84.7270673 |
| 83.1241668 | PD_98059 | 84.9957761 |
| 83.7131655 | Fascaplysin_Synthetic | 85.4585918 |
| 83.7597072 | VEGFRTK_Inhb_II | 87.282199 |
| 83.9387899 | VEGFR2K_Inhb_IV | 87.6244015 |
| 84.0812878 | Bosutinib | 90.3338662 |
| 84.4185614 | SB202190 | 91.0798364 |
| 84.8661647 | MNK1_Inhb | 91.6201253 |
| 86.8288923 | GTP-14564 | 92.4720828 |
| 87.1147601 | Flt-3_Inhb_III | 93.0801954 |
| 88.8037669 | PDGFRTK_Inhb_IV | 94.7169452 |
| 90.4622257 | PKCbII/EGFR_Inhb | 95.6234696 |
| 91.1507508 | GSK3_Inhb_IX | 96.4844973 |
| 93.2100131 | Aurora/Cdk_Inhb | 96.9024234 |
| 93.9422884 | Pazopanib | 97.9234713 |
| 94.4367266 | SU6656 | 101.143278 |
| 94.4660587 | Sunitinib | 101.586093 |
| 94.9556147 | Dovitinib | 107.807139 |
| 101.300549 | SB218078 | 111.923754 |

| | | |
|---|---|---|
| 104.076439 | SU11652 | 113.494017 |
| 109.898583 | PD_169316 | 119.495072 |
| 110.299181 | PDK1/Akt/Flt | 120.042773 |
| 112.235964 | Staurosporine_N | 122.744312 |
| 116.639351 | MetK_Inhb | 124.742623 |
| 118.652719 | H-89_Di | 138.444109 |
| 139.183545 | K-252a | 140.061988 |
| 140.759386 | Cdk1/2_Inhb_III | 140.769883 |
| 145.844986 | TWS119 | 166.74055 |
| 147.266136 | Sorafenib | 189.733312 |

**isolate B**

**se predictions** **drug predi**

| hypnozoites | | Schizonts | |
|---|---|---|---|
| Gene ID | Coefficient | Drug | Prediction |
| AKT1 | -0.5497513 | PKR_Inhb_Neg_Cntr | 21.93368 |
| BRAF | -0.3419016 | VX-702 | 29.54965 |
| CAMK4 | 0.87451717 | VEGFRTK_Inhb_III_KRN633 | 30.45129 |
| CHUK | 0.20006266 | CaseinK_II_Inhb_III_TBCA | 35.58869 |
| CSK | -0.0043528 | Nilotinib | 37.74129 |
| CSNK1A1 | -0.1932317 | DNA-PK_Inhb_III | 38.99001 |
| CSNK2A1 | 0.36171118 | PI_3-Kg_Inhb | 39.19959 |
| DMPK | 0.60594185 | PP3 | 40.08512 |
| DYRK4 | 0.58763392 | PI_3-Kg_Inhb_II | 42.63282 |
| EPHB4 | 0.16590199 | JNK_Inhb_V | 42.98576 |
| ERBB2 | 0.00804346 | AG_1295 | 44.4734 |
| ERBB4 | 0.13491547 | p38_MAPK_Inhb | 45.22197 |
| FLT1 | -0.4574981 | Roscovitine | 46.31099 |
| GRK3 | 0.5522592 | TGF-b_RI_Inhb_III | 46.69358 |
| GRK5 | 0.2064367 | MetK_Inhb | 47.59494 |
| GSK3A | 0.1129771 | PD_169316 | 47.65651 |
| HIPK4 | -0.6863241 | JNK_Inhb_IX | 47.82488 |
| JAK1 | -0.0669944 | Cdk2_Inhb_III | 48.01033 |
| MAP3K14 | -0.3262994 | AG_9 | 48.94367 |
| MST1R | 0.13134244 | Imatinib | 48.96723 |
| NEK11 | -0.0566966 | JAK3_Inhb_IV | 49.31232 |
| NLK | 0.07596804 | DNA-PK_Inhb_II | 49.34533 |
| OXSR1 | 0.38301203 | p38_MAPK_Inhb_III | 49.47951 |
| PAK2 | -0.1146037 | Lapatinib | 49.48392 |
| PIM3 | 0.12466191 | Cdk4_Inhb | 49.4852 |
| PRKACA | -0.2137447 | MNK1_Inhb | 49.6824 |
| PRKCZ | -0.0619058 | LY_303511-_Negative_control | 49.77175 |
| PTK2B | -0.1497714 | PKCbII/EGFR_Inhb | 49.78565 |
| SRPK2 | -0.2975894 | JNK_Inhb_VIII | 50.26175 |
| STK11 | -0.0806712 | GSK3_Inhb_X | 50.37524 |
| STK25 | 0.2032991 | Tandutinib | 50.78603 |
| TAOK2 | -0.0562455 | Alsterpaullone_2Cyanoethyl | 50.92293 |
| TBK1 | 0.01277198 | BAY_11_7082 | 51.20532 |
| TEK | -0.0183146 | JNK_Inhb_Neg_Cntr | 51.30713 |
| TYK2 | -0.0095873 | EGFR_Inhb | 53.20105 |
| TYRO3 | -0.2758903 | MEK_Inhb_I | 53.4135 |
| WNK2 | 0.49379627 | PDGFRTK_Inhb_III | 53.81914 |
| | | AG_490 | 54.25487 |
| | | IC261 | 54.80434 |

| | |
|---|---|
| PI-103 | 55.08009 |
| Bisindolylmaleimide_I | 55.15102 |
| PDGFRTK_Inhb_IV | 55.1693 |
| SB220025 | 55.27333 |
| PKR_Inhb | 55.34696 |
| AG_1296 | 55.42288 |
| ROCK_InhbY | 55.44924 |
| LY294002 | 56.08531 |
| MEK1/2_Inhb | 56.13182 |
| Herbimycin_A_Streptomyces_s | 56.18636 |
| Compound_52 | 56.30883 |
| VEGFR2K_Inhb_IV | 57.31352 |
| Flt-3_Inhb | 57.91576 |
| Dasatinib | 58.357 |
| Fascaplysin_Synthetic | 58.35956 |
| NF-kB_Activation_Inhb | 58.42513 |
| VEGFR2K_Inhb_II | 59.38186 |
| Syk_Inhb_II | 59.43746 |
| Cdk1/5_Inhb | 59.54589 |
| Wortmannin | 59.9278 |
| Mubritinib | 60.44857 |
| Vatalanib | 60.48013 |
| GSK3b_Inhb_XI | 60.55472 |
| JAK_Inhb_I | 61.45906 |
| Cdk2_Inhb_IV_NU6140 | 61.96039 |
| SU9516 | 62.11224 |
| SKF-86002 | 62.14794 |
| GSK3b_Inhb_II | 62.2936 |
| STO-609 | 62.65923 |
| Ro-32-0432 | 62.69651 |
| Flt-3_Inhb_II | 62.8065 |
| VEGFRTK_Inhb_II | 63.87965 |
| Masitinib | 64.15432 |
| Kenpaullone | 64.34189 |
| PP1_Analog_II | 64.48459 |
| PKCb_Inhb | 64.54229 |
| Aurora/Cdk_Inhb | 64.92722 |
| PD_158780 | 64.94474 |
| PDGFRTK_Inhb_II | 65.01907 |
| Akt_Inhb_VIII_Isozyme-Selectiv | 65.16017 |
| Isogranulatimide | 65.37603 |
| Chelerythrine_Chloride | 65.88306 |
| KN-62 | 65.99827 |

| | |
|---|---|
| MK2a_Inhb | 66.03907 |
| Tpl2_K_Inhb | 66.164 |
| Cdk1_Inhb_CGP74514A | 66.38509 |
| SB202474 | 66.65125 |
| IGF-1R_Inhb_II | 66.73293 |
| TGF-b_RI_K_Inhb | 67.04675 |
| MEK_Inhb_II | 67.15199 |
| GSK3_Inhb_XIII | 67.44791 |
| IRAK-1/4_Inhb | 67.56202 |
| Erlotinib | 67.66244 |
| Chk2_Inhb_II | 67.67911 |
| KN-93 | 67.72609 |
| PDK1/Akt/Flt | 67.85432 |
| JNK_Inhb_II | 67.91929 |
| ERK_Inhb_III | 68.55659 |
| Gefitinib | 68.77133 |
| AMPK_Inhb_Comp_C | 68.83173 |
| Diacylglycerol_K_Inhb_II | 69.33997 |
| DNA-PK_Inhb_V | 69.53971 |
| cFMS_RTK_Inhb | 69.89495 |
| AG_112 | 69.90992 |
| Flt-3_Inhb_III | 70.02244 |
| Syk_Inhb_III | 70.1778 |
| SB203580 | 70.24069 |
| Vandetanib | 70.80485 |
| Tofacitinib | 71.38654 |
| Akt_Inhb_X | 71.4449 |
| Rho_K_Inhb_III_Rockout | 71.46811 |
| Purvalanol_A | 71.67373 |
| Cdk4_Inhb_II_NSC_625987 | 71.68661 |
| Akt_Inhb_V_Triciribine | 72.23556 |
| PD_98059 | 72.37969 |
| JAK3_Inhb_VI | 72.69565 |
| Indirubin_E804 | 73.07783 |
| Aminopurvalanol_A | 73.24911 |
| Rapamycin | 73.36732 |
| PD_174265 | 73.72395 |
| SB202190 | 74.13005 |
| G_6983 | 74.16355 |
| Sphingosine_K_Inhb | 74.32211 |
| DMBI | 74.8921 |
| Cdc2-LikeK_Inhb_TG003 | 74.90527 |
| Bohemine | 75.04185 |

| | |
|---|---|
| ERK_Inhb_II_FR180204 | 75.25036 |
| SC-68376 | 75.87981 |
| G_6976 | 75.99276 |
| Tozasertib | 76.05147 |
| AGL_2043 | 76.36093 |
| Bisindolylmaleimide_IV | 76.78145 |
| VEGF_Receptor_2_K_Inhb_I | 77.27704 |
| Indirubin-3,Ä≤-monoxime | 77.29364 |
| Compound_56 | 77.34261 |
| JAK3_Inhb_II | 77.4824 |
| HA_1077_Dihydrochloride_Fasu | 77.66786 |
| GSK3b_Inhb_I | 78.06101 |
| Cdk/Crk_Inhb | 78.74632 |
| Rho_K_Inhb_IV | 79.46894 |
| GSK3b_Inhb_VIII | 79.9581 |
| Akt_Inhb_IV | 80.03859 |
| GTP-14564 | 80.55636 |
| AG_1478 | 80.6194 |
| Staurosporine_N | 80.80652 |
| PDGFRTK_Inhb | 81.79928 |
| Aloisine_RP106 | 82.27705 |
| EGFR/ErbB-2_Inhb | 82.41072 |
| EGFR/ErbB-2/4_Inhb | 82.4594 |
| Bcr-abl_Inhb | 82.92432 |
| Sunitinib | 83.07144 |
| Aurora_Inhb_III | 83.12417 |
| Alsterpaullone | 83.71317 |
| Cdk1_Inhb | 83.75971 |
| Pazopanib | 83.93879 |
| Sorafenib | 84.08129 |
| Cdk4_Inhb_III | 84.41856 |
| CaseinK_I_Inhb_D44 | 84.86616 |
| BPIQ-I | 86.82889 |
| AG_1024 | 87.11476 |
| ATM/ATR_K_Inhb | 88.80377 |
| VEGFR2K_Inhb_III | 90.46223 |
| ATM_K_Inhb | 91.15075 |
| Syk_Inhb | 93.21001 |
| GSK3_Inhb_IX | 93.94229 |
| LCK_Inhb | 94.43673 |
| Aloisine_A_RP107 | 94.46606 |
| Dovitinib | 94.95561 |
| IKK-2_Inhb_IV | 101.30055 |

| | |
|---|---|
| SrcK_Inhb_I | 104.07644 |
| SU11652 | 109.89858 |
| SB218078 | 110.29918 |
| Bosutinib | 112.23596 |
| Cdk1/2_Inhb_III | 116.63935 |
| SU6656 | 118.65272 |
| Staurosporine | 139.18354 |
| H-89_Di | 140.75939 |
| TWS119 | 145.84499 |
| K-252a | 147.26614 |

is

**ictions** | **kinase predictions**

| **Hypnozoites** | | **Schizonts** | |
|---|---|---|---|
| Drug | Prediction | Gene ID | Coefficient |
| JNK_Inhb_V | 0.06309739 | ABL1 | 0.02201779 |
| Staurosporine | 8.49962448 | ABL2 | 0.0257807 |
| Alsterpaullone_2Cyanoethyl | 13.4944128 | AKT1 | 0.05112702 |
| Alsterpaullone | 19.1180133 | BLK | 0.03266699 |
| Bisindolylmaleimide_I | 24.7024328 | BMX | 0.06043793 |
| Aloisine_RP106 | 25.9776593 | CDC42BPA | 1.047676 |
| Bcr-abl_Inhb | 30.8224095 | CDK7 | 0.2737111 |
| AG_1295 | 31.5908324 | CSNK1G2 | -0.0284738 |
| PI_3-Kg_Inhb_II | 32.1415102 | DDR2 | 0.00024851 |
| CaseinK_II_Inhb_III_TBCA | 34.2204541 | DYRK1B | -0.0357072 |
| Tpl2_K_Inhb | 37.6306396 | EPHA7 | -0.0002299 |
| Cdk1/5_Inhb | 39.2480951 | FLT1 | -0.1098778 |
| Cdk4_Inhb | 40.1166801 | FRK | -0.1009732 |
| Dasatinib | 40.8491277 | GRK3 | -1.540287 |
| TGF-b_RI_K_Inhb | 41.7767675 | HIPK2 | -0.6979694 |
| Erlotinib | 42.0171284 | HIPK3 | 0.2670111 |
| Gefitinib | 43.5088272 | HIPK4 | -0.4898929 |
| Bisindolylmaleimide_IV | 44.5194687 | JAK1 | -0.1362244 |
| GSK3_Inhb_XIII | 44.7178411 | JAK2 | -0.246369 |
| Aloisine_A_RP107 | 44.84552 | MAP3K7 | -0.0967505 |
| Akt_Inhb_X | 45.1166186 | MAP4K5 | 0.107783 |
| ATM_K_Inhb | 45.8273083 | MAPK1 | -0.0043671 |
| AG_490 | 46.0395709 | MAPK8 | 0.1333497 |
| GSK3_Inhb_X | 46.4935455 | MAPK9 | 0.2573735 |
| BAY_11_7082 | 46.6529186 | MAPKAPK5 | 0.0327382 |
| JAK3_Inhb_IV | 47.7205023 | MERTK | 0.00459151 |
| ROCK_InhbY | 48.0507303 | MKNK1 | -0.2627957 |
| ATM/ATR_K_Inhb | 48.3627591 | MST1R | 1.145722 |
| Roscovitine | 48.5488074 | NEK2 | 0.4048569 |
| Compound_56 | 48.7000839 | NEK7 | -0.2762704 |
| Aurora_Inhb_III | 49.5396353 | NUAK1 | 0.00609363 |
| GSK3b_Inhb_XI | 50.3151276 | OXSR1 | -0.0549601 |
| DNA-PK_Inhb_II | 50.5011752 | PAK5 | -0.2766588 |
| Akt_Inhb_IV | 50.6298318 | PBK | -0.7939815 |
| AMPK_Inhb_Comp_C | 51.2631313 | PLK2 | -0.0691325 |
| Rho_K_Inhb_IV | 51.4730317 | PNCK | -0.1154872 |
| Cdk1_Inhb_CGP74514A | 52.1997141 | RAF1 | 0.09913362 |
| PKCb_Inhb | 52.3579363 | TEK | -0.4550193 |
| Tandutinib | 52.6448527 | TYK2 | -0.0103729 |

| | |
|---|---|
| PKR_Inhb | 52.6686287 |
| JNK_Inhb_VIII | 53.1149832 |
| Cdc2-LikeK_Inhb_TG003 | 53.5009139 |
| STO-609 | 53.6326193 |
| VX-702 | 53.6753273 |
| PI_3-Kg_Inhb | 53.6756323 |
| NF-kB_Activation_Inhb | 55.3356951 |
| Akt_Inhb_VIII_Isozyme-Selective_A | 55.3750906 |
| LY_303511-_Negative_control | 55.6052176 |
| Cdk2_Inhb_IV_NU6140 | 56.2601119 |
| p38_MAPK_Inhb | 56.6065027 |
| GSK3b_Inhb_I | 57.3130805 |
| p38_MAPK_Inhb_III | 57.3515645 |
| Cdk/Crk_Inhb | 57.5672723 |
| Ro-32-0432 | 57.6287981 |
| Bohemine | 57.7131497 |
| Lapatinib | 57.7690389 |
| AG_1024 | 57.8919277 |
| Flt-3_Inhb | 58.1914854 |
| Cdk1_Inhb | 58.9286296 |
| VEGFR2K_Inhb_II | 59.065591 |
| Sphingosine_K_Inhb | 59.3304089 |
| Mubritinib | 59.4227344 |
| Cdk4_Inhb_II_NSC_625987 | 59.5304789 |
| AG_112 | 60.3122065 |
| Kenpaullone | 60.4278262 |
| AGL_2043 | 60.6413845 |
| Isogranulatimide | 60.7937243 |
| Chelerythrine_Chloride | 61.0252852 |
| DNA-PK_Inhb_V | 61.3883119 |
| JNK_Inhb_Neg_Cntr | 61.4344367 |
| TGF-b_RI_Inhb_III | 61.54287 |
| Chk2_Inhb_II | 61.6623911 |
| GSK3b_Inhb_VIII | 61.9759168 |
| DNA-PK_Inhb_III | 62.2092207 |
| GSK3b_Inhb_II | 62.7361537 |
| Cdk4_Inhb_III | 62.8640495 |
| CaseinK_I_Inhb_D44 | 62.9772508 |
| LY294002 | 63.1500396 |
| Masitinib | 63.3351803 |
| Syk_Inhb_III | 63.5803401 |
| ERK_Inhb_II_FR180204 | 63.8461473 |
| KN-93 | 63.9344906 |

| | |
|---|---|
| Wortmannin | 64.0726025 |
| SU9516 | 64.3351865 |
| EGFR/ErbB-2_Inhb | 64.4886396 |
| ERK_Inhb_III | 64.7054802 |
| Akt_Inhb_V_Triciribine | 64.8019045 |
| Imatinib | 65.0125395 |
| JNK_Inhb_II | 66.2186294 |
| Indirubin_E804 | 66.3564885 |
| EGFR/ErbB-2/4_Inhb | 66.8642041 |
| VEGF_Receptor_2_K_Inhb_I | 67.1020951 |
| Tozasertib | 67.1822257 |
| Rho_K_Inhb_III_Rockout | 67.9707921 |
| PKR_Inhb_Neg_Cntr | 68.0918683 |
| MEK_Inhb_II | 68.2828156 |
| SrcK_Inhb_I | 68.7139422 |
| JAK_Inhb_I | 69.1341107 |
| Syk_Inhb_II | 69.5378602 |
| SB202474 | 69.5604833 |
| IGF-1R_Inhb_II | 70.0915612 |
| DMBI | 70.092429 |
| BPIQ-I | 70.0985554 |
| Syk_Inhb | 70.1198943 |
| HA_1077_Dihydrochloride_Fasudil | 70.1758344 |
| Herbimycin_A_Streptomyces_sp. | 70.4609487 |
| PD_158780 | 70.6193677 |
| Purvalanol_A | 71.0628471 |
| Compound_52 | 71.2070597 |
| SB203580 | 71.4241107 |
| Vatalanib | 71.4249744 |
| cFMS_RTK_Inhb | 72.0403715 |
| G_6976 | 72.0839909 |
| JAK3_Inhb_VI | 72.1020885 |
| Cdk2_Inhb_III | 72.2749201 |
| Rapamycin | 72.3480595 |
| AG_9 | 72.6353418 |
| JNK_Inhb_IX | 72.674416 |
| Nilotinib | 72.8087342 |
| Vandetanib | 72.859401 |
| G_6983 | 72.9501463 |
| IC261 | 73.5865134 |
| Flt-3_Inhb_II | 73.8728023 |
| PDGFRTK_Inhb_II | 74.7052218 |
| LCK_Inhb | 74.7524262 |

| | |
|---|---|
| JAK3_Inhb_II | 74.7756938 |
| Diacylglycerol_K_Inhb_II | 74.7773458 |
| MEK1/2_Inhb | 75.4767097 |
| KN-62 | 75.6636006 |
| Tofacitinib | 76.3492307 |
| SC-68376 | 76.3600781 |
| Indirubin-3,Ä≤-monoxime | 76.8476684 |
| AG_1296 | 77.0242988 |
| SB220025 | 77.4903048 |
| PP3 | 78.6631485 |
| PI-103 | 78.8427986 |
| MEK_Inhb_I | 78.9149695 |
| PP1_Analog_II | 79.3508386 |
| PDGFRTK_Inhb | 79.7385078 |
| SKF-86002 | 80.2140604 |
| VEGFR2K_Inhb_III | 81.5469691 |
| Aminopurvalanol_A | 81.6409521 |
| AG_1478 | 81.6625421 |
| EGFR_Inhb | 81.8942131 |
| VEGFRTK_Inhb_III_KRN633 | 82.0428312 |
| IKK-2_Inhb_IV | 82.7825729 |
| PD_174265 | 83.2994283 |
| IRAK-1/4_Inhb | 84.133721 |
| MK2a_Inhb | 84.5438316 |
| PDGFRTK_Inhb_III | 84.7270673 |
| PD_98059 | 84.9957761 |
| Fascaplysin_Synthetic | 85.4585918 |
| VEGFRTK_Inhb_II | 87.282199 |
| VEGFR2K_Inhb_IV | 87.6244015 |
| Bosutinib | 90.3338662 |
| SB202190 | 91.0798364 |
| MNK1_Inhb | 91.6201253 |
| GTP-14564 | 92.4720828 |
| Flt-3_Inhb_III | 93.0801954 |
| PDGFRTK_Inhb_IV | 94.7169452 |
| PKCbII/EGFR_Inhb | 95.6234696 |
| GSK3_Inhb_IX | 96.4844973 |
| Aurora/Cdk_Inhb | 96.9024234 |
| Pazopanib | 97.9234713 |
| SU6656 | 101.143278 |
| Sunitinib | 101.586093 |
| Dovitinib | 107.807139 |
| SB218078 | 111.923754 |

| | |
|---|---|
| SU11652 | 113.494017 |
| PD_169316 | 119.495072 |
| PDK1/Akt/Flt | 120.042773 |
| Staurosporine_N | 122.744312 |
| MetK_Inhb | 124.742623 |
| H-89_Di | 138.444109 |
| K-252a | 140.061988 |
| Cdk1/2_Inhb_III | 140.769883 |
| TWS119 | 166.74055 |
| Sorafenib | 189.733311 |

**olate C**

**drug predictions**

| Schizonts | |
|---|---|
| Drug | Prediction |
| PD_169316 | 15.53096 |
| JNK_Inhb_VIII | 32.50538 |
| JNK_Inhb_IX | 44.584 |
| GTP-14564 | 47.7748 |
| KN-62 | 60.65764 |
| GSK3_Inhb_IX | 60.97775 |
| Kenpaullone | 66.63102 |
| Imatinib | 67.35866 |
| EGFR_Inhb | 72.54795 |
| ERK_Inhb_II_FR180204 | 72.57842 |
| H-89_Di | 73.77626 |
| p38_MAPK_Inhb_III | 74.55505 |
| ERK_Inhb_III | 75.48819 |
| IC261 | 75.76669 |
| GSK3b_Inhb_XI | 76.20731 |
| GSK3b_Inhb_VIII | 76.60663 |
| KN-93 | 78.07976 |
| GSK3_Inhb_X | 78.87517 |
| MEK_Inhb_II | 79.9513 |
| PI-103 | 81.17221 |
| MetK_Inhb | 81.1807 |
| Flt-3_Inhb_II | 82.61833 |
| Flt-3_Inhb | 82.73329 |
| PI_3-Kg_Inhb | 84.00039 |
| IGF-1R_Inhb_II | 85.02055 |
| MEK_Inhb_I | 85.06713 |
| Vatalanib | 85.35591 |
| LY_303511-_Negative_con | 85.53282 |
| DNA-PK_Inhb_V | 85.6122 |
| PKR_Inhb_Neg_Cntr | 85.66976 |
| Tpl2_K_Inhb | 86.52195 |
| EGFR/ErbB-2/4_Inhb | 87.45795 |
| DNA-PK_Inhb_II | 88.69064 |
| G_6983 | 88.86578 |
| Cdk4_Inhb_II_NSC_625987 | 89.13241 |
| Pazopanib | 89.84669 |
| SC-68376 | 90.55761 |
| EGFR/ErbB-2_Inhb | 91.11514 |
| Indirubin-3,Ä≤-monoxime | 91.43682 |

| | |
|---|---|
| Lapatinib | 91.87544 |
| VX-702 | 92.43191 |
| Syk_Inhb | 93.81679 |
| NF-kB_Activation_Inhb | 94.15379 |
| ATM/ATR_K_Inhb | 94.52564 |
| HA_1077_Dihydrochloride_ | 94.60745 |
| Diacylglycerol_K_Inhb_II | 94.88419 |
| cFMS_RTK_Inhb | 94.8992 |
| VEGFR2K_Inhb_II | 95.21264 |
| SB202190 | 95.39137 |
| Wortmannin | 95.39799 |
| JNK_Inhb_V | 95.6551 |
| PD_98059 | 96.11525 |
| MEK1/2_Inhb | 96.22713 |
| Sphingosine_K_Inhb | 96.65399 |
| JAK3_Inhb_IV | 96.89815 |
| Herbimycin_A_Streptomyc | 97.48667 |
| SU9516 | 97.69714 |
| PDGFRTK_Inhb | 97.9677 |
| Bosutinib | 98.43364 |
| PP1_Analog_II | 98.44784 |
| Akt_Inhb_IV | 98.51272 |
| LCK_Inhb | 98.74548 |
| AG_1295 | 98.95523 |
| Bisindolylmaleimide_IV | 99.7387 |
| Tozasertib | 99.81369 |
| PD_158780 | 99.91228 |
| Alsterpaullone_2Cyanoeth | 100.20215 |
| PKCbII/EGFR_Inhb | 100.22232 |
| Nilotinib | 100.39093 |
| Bohemine | 100.79469 |
| AG_1024 | 100.94446 |
| JAK3_Inhb_II | 101.06473 |
| VEGFR2K_Inhb_IV | 101.09064 |
| MNK1_Inhb | 101.29919 |
| VEGF_Receptor_2_K_Inhb_ | 101.38816 |
| PDGFRTK_Inhb_II | 101.78184 |
| GSK3b_Inhb_I | 102.55399 |
| JNK_Inhb_Neg_Cntr | 102.65694 |
| SB203580 | 102.76484 |
| LY294002 | 102.95665 |
| TGF-b_RI_Inhb_III | 104.05152 |
| PP3 | 104.10131 |

| | |
|---|---|
| Cdk/Crk_Inhb | 104.27996 |
| Flt-3_Inhb_III | 104.3789 |
| Erlotinib | 105.29721 |
| DNA-PK_Inhb_III | 105.43573 |
| p38_MAPK_Inhb | 105.51084 |
| PDGFRTK_Inhb_III | 106.2077 |
| Ro-32-0432 | 106.27435 |
| IRAK-1/4_Inhb | 106.31266 |
| DMBI | 106.6431 |
| SU6656 | 107.11695 |
| Roscovitine | 107.93385 |
| PI_3-Kg_Inhb_II | 108.4413 |
| JNK_Inhb_II | 108.89505 |
| SB202474 | 109.01576 |
| Purvalanol_A | 109.50885 |
| GSK3_Inhb_XIII | 109.64736 |
| SB220025 | 109.91853 |
| Gefitinib | 110.46624 |
| MK2a_Inhb | 111.18755 |
| Dasatinib | 111.72957 |
| VEGFRTK_Inhb_II | 112.42438 |
| Rapamycin | 112.71972 |
| Cdk4_Inhb_III | 112.76499 |
| Compound_52 | 112.82766 |
| SKF-86002 | 112.88598 |
| Aurora_Inhb_III | 113.03109 |
| Compound_56 | 113.57406 |
| Indirubin_E804 | 113.73583 |
| Cdk1_Inhb | 114.12558 |
| Fascaplysin_Synthetic | 114.5273 |
| Chelerythrine_Chloride | 114.55415 |
| Cdk2_Inhb_III | 114.75275 |
| AGL_2043 | 115.58735 |
| PD_174265 | 115.80445 |
| Cdc2-LikeK_Inhb_TG003 | 116.63639 |
| AG_112 | 116.75881 |
| VEGFR2K_Inhb_III | 116.89111 |
| Isogranulatimide | 117.00452 |
| ROCK_InhbY | 117.80376 |
| Vandetanib | 117.94324 |
| VEGFRTK_Inhb_III_KRN633 | 119.05769 |
| STO-609 | 119.40627 |
| Cdk1/5_Inhb | 119.42137 |

| | |
|---|---|
| Cdk4_Inhb | 119.61509 |
| PKCb_Inhb | 119.68632 |
| CaseinK_I_Inhb_D44 | 119.9666 |
| Akt_Inhb_V_Triciribine | 120.04911 |
| ATM_K_Inhb | 120.08155 |
| CaseinK_II_Inhb_III_TBCA | 120.18879 |
| Cdk2_Inhb_IV_NU6140 | 121.48424 |
| Aloisine_RP106 | 121.76867 |
| SrcK_Inhb_I | 121.80203 |
| AG_1296 | 121.88081 |
| Chk2_Inhb_II | 123.21045 |
| Cdk1_Inhb_CGP74514A | 123.26172 |
| Mubritinib | 124.92646 |
| Rho_K_Inhb_IV | 125.23299 |
| TGF-b_RI_K_Inhb | 125.29637 |
| BPIQ-I | 125.82163 |
| TWS119 | 126.05154 |
| AG_1478 | 126.09064 |
| Akt_Inhb_VIII_Isozyme-Sel | 127.95494 |
| Akt_Inhb_X | 129.34337 |
| GSK3b_Inhb_II | 131.1183 |
| IKK-2_Inhb_IV | 131.99044 |
| Rho_K_Inhb_III_Rockout | 132.49719 |
| PKR_Inhb | 132.67695 |
| Syk_Inhb_III | 132.7129 |
| G_6976 | 133.09652 |
| PDGFRTK_Inhb_IV | 134.30692 |
| Tandutinib | 135.14673 |
| Alsterpaullone | 135.26947 |
| Masitinib | 137.17304 |
| Sorafenib | 139.32242 |
| BAY_11_7082 | 140.00534 |
| Sunitinib | 140.87459 |
| AG_490 | 141.31838 |
| AG_9 | 144.12607 |
| Dovitinib | 144.20585 |
| Staurosporine_N | 144.81884 |
| JAK3_Inhb_VI | 144.9709 |
| Aloisine_A_RP107 | 147.7898 |
| Bisindolylmaleimide_I | 150.1836 |
| SU11652 | 155.64524 |
| PDK1/Akt/Flt | 156.77666 |
| AMPK_Inhb_Comp_C | 158.36402 |

| | |
|---|---|
| JAK_Inhb_I | 163.81999 |
| Aminopurvalanol_A | 169.91977 |
| Syk_Inhb_II | 172.90401 |
| Cdk1/2_Inhb_III | 175.83038 |
| Bcr-abl_Inhb | 177.52376 |
| Tofacitinib | 179.0799 |
| Aurora/Cdk_Inhb | 212.18739 |
| Staurosporine | 225.69877 |
| K-252a | 237.30691 |
| SB218078 | 250.58688 |
