## Supplementary File 1 - Raw data for "Host kinase regulation of *Plasmodium vivax* dormant and replicating liver stages"

**Parasites/well**

| isolate A | isolate B | isolate C |
|---|---|---|
| 52 | 52 | 24 |
| 52 | 31 | 15 |
| 69 | 45 | 18 |
| 31 | 26 | 14 |
| 31 | 33 | 16 |
| 21 | 39 | 33 |
| 20 | 36 | |
| 25 | 30 | |
| 20 | 62 | |
| 39 | | |
| 28 | | |
| 73 | | |
| 46 | | |
| 40 | | |
| 31 | | |
| 25 | | |
| 24 | | |
| 42 | | |

**hypnozoites:schizonts**

| isolate A | isolate B | isolate C |
|---|---|---|
| 0.7931035 | 0.1555556 | 0.2 |
| 0.9259259 | 0.2916667 | 0.0714286 |
| 0.5681818 | 0.1842105 | 0.0588235 |
| 0.7222222 | 0.4444444 | 0.0769231 |
| 1.2142857 | 0.137931 | 0.1428571 |
| 1.625 | 0.56 | 0.137931 |
| 1.2222222 | 0.44 | |
| 0.9230769 | 0.7647059 | |
| 1.8571429 | 0.24 | |
| 0.4444444 | | |
| 0.5555556 | | |
| 0.46 | | |
| 0.6428571 | | |
| 0.3793103 | | |
| 0.8235294 | | |
| 1.5 | | |
| 2 | | |
| 0.4482759 | | |

**schizont size**

| isolate A | isolate B | isolate C |
|---|---|---|
| 904.35 | 240.81 | 405.37 |
| 666.56 | 668.07 | 1028.15 |
| 1163.27 | 474.06 | 2120.46 |
| 984.36 | 425.75 | 625.8 |
| 614.47 | 245.34 | 552.57 |
| 554.08 | 428.77 | 709.59 |
| 430.28 | 208.35 | 917.93 |
| 248.36 | 352.53 | 619.76 |
| 493.69 | 202.31 | 496.71 |
| 252.88 | 578.99 | 730.72 |
| 441.6 | 542 | 805.46 |
| 532.19 | 341.96 | 2161.98 |
| 225.71 | 674.86 | 166.83 |
| 485.39 | 753.37 | 875.66 |
| 515.58 | 1234.23 | 1798.12 |
| 184.19 | 873.4 | 190.98 |
| 420.47 | 1041.73 | 294.4 |
| 557.1 | 631.08 | 403.86 |
| 317.8 | 298.93 | 1424.46 |
| 337.43 | 197.02 | 1493.15 |
| 626.55 | 542 | 637.12 |
| 379.7 | 1706.03 | 1359.54 |
| 736.76 | 720.16 | 1807.94 |
| 768.47 | 258.92 | 1398.04 |
| 575.97 | 474.82 | 708.08 |
| 664.29 | 447.64 | 1496.93 |
| 723.93 | 421.22 | 1846.44 |
| 694.49 | 1133.83 | 1400.3 |
| 398.58 | | |

**Isolate A**

Schizont counts

| | | | | | | |
|---|---|---|---|---|---|---|
| DMSO | 29 | 27 | 44 | 18 | 14 | 8 |
| KI1 | 19 | 14 | 9 | | | |
| KI2 | 31 | 21 | 17 | | | |
| KI3 | 15 | 11 | 15 | | | |
| KI4 | 30 | 21 | 7 | | | |
| KI5 | 25 | 19 | 6 | | | |
| KI6 | 6 | 5 | 8 | | | |
| KI7 | 13 | 29 | 5 | | | |
| KI8 | 17 | 15 | 23 | | | |
| KI10 | 4 | 10 | 7 | | | |
| KI11 | 12 | 9 | 10 | | | |
| KI13 | 6 | 32 | 3 | | | |
| KI14 | 18 | 16 | 25 | | | |
| KI15 | 15 | 5 | 11 | | | |
| KI16 | 5 | 19 | 6 | | | |
| KI17 | 15 | 19 | 17 | | | |
| KI18 | 26 | 5 | 10 | | | |
| KI19 | 20 | 17 | 23 | | | |
| KI20 | 40 | 39 | 26 | | | |
| KI21 | 16 | 5 | 6 | | | |
| KI22 | 14 | 6 | 6 | | | |
| KI23 | 12 | 29 | 11 | | | |
| KI25 | 5 | 10 | 1 | | | |
| KI26 | 11 | 10 | 16 | | | |
| KI27 | 28 | 7 | 10 | | | |
| KI28 | 8 | 7 | 4 | | | |
| KI30 | 14 | 20 | 25 | | | |
| KI31 | 2 | 2 | 3 | | | |
| KI32 | 16 | 9 | 11 | | | |
| KI33 | 14 | 10 | 10 | | | |
| KI34 | 10 | 20 | 10 | | | |
| KI35 | 8 | 7 | 9 | | | |
| KI36 | 21 | 9 | 27 | | | |
| KI37 | 16 | 4 | 11 | | | |
| KI38 | 16 | | 25 | | | |

Hypnozoite counts

| | | | | | | |
|---|---|---|---|---|---|---|
| DMSO | 23 | 25 | 25 | 13 | 17 | 13 |
| KI1 | 15 | 20 | 5 | | | |
| KI2 | 15 | 19 | 15 | | | |
| KI3 | 12 | 14 | 7 | | | |
| KI4 | 22 | 31 | 14 | | | |

| | | | |
|---|---|---|---|
| KI5 | 19 | 14 | 11 |
| KI6 | 5 | 4 | 12 |
| KI7 | 12 | 26 | 9 |
| KI8 | 10 | 6 | 13 |
| KI10 | 6 | 20 | 8 |
| KI11 | 16 | 8 | 6 |
| KI13 | 15 | 45 | 11 |
| KI14 | 13 | 16 | 22 |
| KI15 | 13 | 9 | 13 |
| KI16 | 14 | 19 | 17 |
| KI17 | 22 | 16 | 23 |
| KI18 | 34 | 9 | 15 |
| KI19 | 30 | 31 | 35 |
| KI20 | 36 | 43 | 26 |
| KI21 | 33 | 11 | 5 |
| KI22 | 17 | 13 | 8 |
| KI23 | 20 | 16 | 18 |
| KI25 | 21 | 26 | 12 |
| KI26 | 21 | 19 | 12 |
| KI27 | 22 | 11 | 14 |
| KI28 | 20 | 21 | 5 |
| KI30 | 19 | 17 | 32 |
| KI31 | 18 | 11 | 14 |
| KI32 | 13 | 20 | 14 |
| KI33 | 18 | 8 | 21 |
| KI34 | 18 | 28 | 16 |
| KI35 | 14 | 16 | 16 |
| KI36 | 17 | 12 | 15 |
| KI37 | 27 | 14 | 22 |
| KI38 | 18 | | 11 |

| | | | | | | |
|---|---|---|---|---|---|---|
| 9 | 13 | 7 | 27 | 18 | 50 | 28 |
| 11 | 12 | 13 | 12 | 10 | 23 | 18 |

**Isolate B**

| | | | | | | |
|---|---|---|---|---|---|---|
| | | | | | Schizont counts | |
| 29 | 17 | 10 | 8 | 29 | DMSO | 45 |
| | | | | | KI1 | 22 |
| | | | | | KI2 | 26 |
| | | | | | KI3 | 21 |
| | | | | | KI4 | 21 |
| | | | | | KI5 | 11 |
| | | | | | KI6 | 35 |
| | | | | | KI7 | 7 |
| | | | | | KI8 | 22 |
| | | | | | KI10 | 23 |
| | | | | | KI11 | 12 |
| | | | | | KI13 | 26 |
| | | | | | KI14 | 28 |
| | | | | | KI15 | |
| | | | | | KI16 | 16 |
| | | | | | KI17 | 67 |
| | | | | | KI18 | 12 |
| | | | | | KI19 | 23 |
| | | | | | KI20 | 40 |
| | | | | | KI21 | 13 |
| | | | | | KI22 | 27 |
| | | | | | KI23 | 13 |
| | | | | | KI25 | 17 |
| | | | | | KI26 | 46 |
| | | | | | KI27 | 16 |
| | | | | | KI28 | 21 |
| | | | | | KI30 | 20 |
| | | | | | KI31 | 13 |
| | | | | | KI32 | 30 |
| | | | | | KI33 | 29 |
| | | | | | KI34 | 26 |
| | | | | | KI35 | 78 |
| | | | | | KI36 | 14 |
| | | | | | KI37 | 8 |
| | | | | | KI38 | 28 |
| | | | | | Hypnozoite counts | |
| 11 | 14 | 15 | 16 | 13 | DMSO | 7 |
| | | | | | KI1 | 4 |
| | | | | | KI2 | 9 |
| | | | | | KI3 | 2 |
| | | | | | KI4 | 4 |

| | |
|---|---|
| KI5 | 1 |
| KI6 | 8 |
| KI7 | 3 |
| KI8 | 7 |
| KI10 | 6 |
| KI11 | 1 |
| KI13 | 4 |
| KI14 | 9 |
| KI15 | |
| KI16 | 6 |
| KI17 | 14 |
| KI18 | 6 |
| KI19 | 9 |
| KI20 | 14 |
| KI21 | 7 |
| KI22 | 9 |
| KI23 | 3 |
| KI25 | 3 |
| KI26 | 9 |
| KI27 | 4 |
| KI28 | 9 |
| KI30 | 10 |
| KI31 | 4 |
| KI32 | 9 |
| KI33 | 3 |
| KI34 | 9 |
| KI35 | 21 |
| KI36 | 7 |
| KI37 | 4 |
| KI38 | 12 |

| | | | | | | |
|---|---|---|---|---|---|---|
| 24 | 38 | 18 | 29 | 25 | 25 | 17 |
| 23 | 21 | | | | | |
| 25 | 35 | | | | | |
| 56 | 25 | | | | | |
| 19 | 16 | | | | | |
| 18 | 16 | | | | | |
| 16 | 26 | | | | | |
| 23 | 21 | | | | | |
| 19 | 21 | | | | | |
| 20 | 32 | | | | | |
| 34 | 16 | | | | | |
| 20 | 23 | | | | | |
| 14 | 25 | | | | | |
| 18 | 11 | | | | | |
| 10 | 34 | | | | | |
| 35 | 27 | | | | | |
| 11 | 20 | | | | | |
| 16 | 16 | | | | | |
| 41 | 53 | | | | | |
| 16 | 13 | | | | | |
| 25 | 34 | | | | | |
| 22 | 24 | | | | | |
| 7 | 26 | | | | | |
| 25 | 28 | | | | | |
| 33 | 28 | | | | | |
| 16 | 24 | | | | | |
| 32 | 20 | | | | | |
| 12 | 16 | | | | | |
| 30 | 50 | | | | | |
| 24 | 18 | | | | | |
| 17 | 23 | | | | | |
| 29 | 26 | | | | | |
| 25 | 25 | | | | | |
| 15 | 25 | | | | | |
| 29 | 48 | | | | | |

| | | | | | | |
|---|---|---|---|---|---|---|
| 7 | 7 | 8 | 4 | 14 | 11 | 13 |
| 11 | 8 | | | | | |
| 10 | 8 | | | | | |
| 16 | 7 | | | | | |
| 7 | 4 | | | | | |

| | |
|---|---|
| 4 | 6 |
| 1 | 8 |
| 6 | 2 |
| 4 | 3 |
| 6 | 6 |
| 7 | 3 |
| 5 | 11 |
| 5 | 6 |
| 3 | 5 |
| 2 | 4 |
| 11 | 14 |
| 8 | 4 |
| 7 | 3 |
| 12 | 13 |
| 4 | 5 |
| 5 | 7 |
| 6 | 8 |
| 4 | 7 |
| 7 | 15 |
| 14 | 35 |
| 1 | 9 |
| 12 | 12 |
| 13 | 18 |
| 8 | 11 |
| 10 | 3 |
| 8 | 5 |
| 16 | 10 |
| 5 | 8 |
| 8 | 1 |
| 12 | 15 |

**Isolate C**

<u>schizont counts</u>

| | | | | | |
|---|---|---|---|---|---|
| 50 | DMSO | 9 | 15 | 3 | 8 |
| | KI1 | 18 | 7 | 14 | |
| | KI2 | 5 | 4 | 4 | |
| | KI3 | 7 | 4 | 11 | |
| | KI4 | 9 | 8 | 10 | |
| | KI5 | 9 | 10 | 8 | |
| | KI6 | 4 | 16 | 7 | |
| | KI7 | 7 | 7 | 11 | |
| | KI8 | 10 | 15 | 11 | |
| | KI10 | 11 | 5 | 3 | |
| | KI11 | 12 | 7 | 5 | |
| | KI13 | 13 | 5 | 12 | |
| | KI14 | 3 | 6 | 11 | |
| | KI15 | 5 | 8 | 5 | |
| | KI16 | 9 | 11 | 5 | |
| | KI17 | 7 | 6 | 3 | |
| | KI18 | 4 | 4 | 5 | |
| | KI19 | 16 | 10 | 11 | |
| | KI20 | 18 | 21 | 15 | |
| | KI21 | 5 | 9 | 7 | |
| | KI22 | 6 | 8 | 9 | |
| | KI23 | 15 | 7 | 10 | |
| | KI25 | 12 | 8 | 10 | |
| | KI26 | 15 | 24 | 18 | |
| | KI27 | 10 | 10 | 12 | |
| | KI28 | 11 | 4 | 10 | |
| | KI30 | 9 | 14 | 10 | |
| | KI31 | 7 | 4 | 7 | |
| | KI32 | 10 | 7 | 7 | |
| | KI33 | 8 | 6 | 10 | |
| | KI34 | 16 | 18 | 8 | |
| | KI35 | 13 | 7 | 9 | |
| | KI36 | 10 | 10 | 7 | |
| | KI37 | 12 | 8 | 7 | |
| | KI38 | 11 | 23 | 6 | |

| | | | | |
|---|---|---|---|---|
| 5 | 10 | 6 | 5 | 7 |

schizonts/well

| | | | |
|---|---|---|---|
| | **SB203580** | | |
| NT (DMSO) | 38 | 45 | 48 |
| 100nM | 27 | 45 | 39 |
| 500nM | 28 | 29 | 34 |
| 1uM | 31 | 29 | 26 |
| 10uM | 13 | 20 | 13 |
| | **JNK inhib V** | | |
| NT (DMSO) | 35 | 23 | 29 |
| 100nM | 43 | 29 | 27 |
| 500nM | 35 | 33 | 31 |
| 1uM | 42 | 41 | 43 |
| 10uM | 18 | 10 | 12 |
| | **PKR inhibitor** | | |
| NT (DMSO) | 33 | 53 | 32 |
| 100nM | 38 | 32 | 31 |
| 500nM | 31 | 49 | 28 |
| 1uM | 41 | 23 | 19 |
| 10uM | 29 | 15 | 27 |
| | **PD169316** | | |
| NT (DMSO) | 45 | 25 | 43 |
| 100nM | 27 | 47 | 61 |
| 500nM | 24 | 40 | 33 |
| 1uM | 22 | 31 | 24 |
| 10uM | 25 | 30 | 30 |
| | **p38 MAPK inhib** | | |
| NT (DMSO) | 25 | 43 | 20 |
| 100nM | 24 | 25 | 43 |
| 500nM | 27 | 42 | 29 |
| 1uM | 34 | 23 | 27 |
| 10uM | 15 | 19 | 31 |

Isolate A - schizont area (µm2)

| DMSO | KI1 | KI2 | KI3 | KI4 | KI5 | KI6 |
|---|---|---|---|---|---|---|
| 447.64384 | 357.05824 | 434.056 | 249.1104 | 132.85888 | 96.62464 | 209.85664 |
| 586.54176 | 512.56352 | 184.19072 | 226.464 | 357.81312 | 950.39392 | 280.81536 |
| 973.04032 | 325.35328 | 468.78048 | 468.78048 | 361.58752 | 488.40736 | 75.488 |
| 658.25536 | 480.10368 | 485.38784 | 220.42496 | 213.63104 | 437.8304 | 160.78944 |
| 1163.2701 | 232.50304 | 392.5376 | 373.6656 | 113.98688 | 329.12768 | 234.0128 |
| 703.54816 | 255.90432 | 326.86304 | 573.7088 | 302.70688 | 273.26656 | |
| 974.55008 | 744.31168 | 357.81312 | 1590.5322 | 351.77408 | 163.80896 | |
| 230.2384 | 757.14464 | 518.60256 | 573.7088 | 676.37248 | 911.14016 | |
| 445.3792 | 517.0928 | 1306.6973 | 531.43552 | 320.824 | 187.96512 | |
| 234.76768 | 305.7264 | 600.1296 | 304.21664 | 285.34464 | 797.90816 | |
| 488.40736 | 543.5136 | 497.46592 | 200.0432 | 304.97152 | 749.59584 | |
| 477.83904 | 502.75008 | 123.04544 | 214.38592 | | 847.73024 | |
| 215.89568 | 606.92352 | 144.93696 | 256.6592 | | 163.05408 | |
| 413.67424 | 1186.6714 | 366.87168 | 443.86944 | | 98.1344 | |
| 513.3184 | 198.53344 | 187.96512 | | | 372.91072 | |
| 190.98464 | | 825.08384 | | | | |
| 280.06048 | | 287.60928 | | | | |
| 542.75872 | | 243.07136 | | | | |
| 454.43776 | | 214.38592 | | | | |
| 318.55936 | | 117.76128 | | | | |
| 600.1296 | | 554.08192 | | | | |
| 369.13632 | | | | | | |
| 722.42016 | | | | | | |
| 714.11648 | | | | | | |
| 553.32704 | | | | | | |
| 599.37472 | | | | | | |
| 701.28352 | | | | | | |
| 641.648 | | | | | | |
| 397.82176 | | | | | | |

| KI7 | KI8 | KI10 | KI11 | KI13 | KI14 | KI15 |
|---|---|---|---|---|---|---|
| 157.76992 | 218.16032 | 203.8176 | 167.58336 | 181.1712 | 328.3728 | 713.3616 |
| 474.81952 | 152.48576 | 545.02336 | 123.80032 | 157.01504 | 254.39456 | 361.58752 |
| 118.51616 | 109.4576 | 98.1344 | 119.27104 | 298.1776 | 350.26432 | 261.18848 |
| 304.97152 | 238.54208 | 144.93696 | 98.1344 | 380.45952 | 358.568 | 1317.2656 |
| 332.1472 | 138.14304 | | 98.1344 | 320.824 | 247.60064 | 359.32288 |
| 273.26656 | 155.50528 | | 134.36864 | 323.08864 | 272.51168 | 196.2688 |
| 249.86528 | 94.36 | | 247.60064 | 255.14944 | 217.40544 | |
| 145.69184 | 224.95424 | | 326.86304 | 258.92384 | 491.42688 | |
| 135.8784 | 88.32096 | | 271.7568 | 748.08608 | 91.34048 | |
| 380.45952 | 146.44672 | | 212.87616 | 452.17312 | 160.03456 | |
| 190.98464 | 301.952 | | 398.57664 | 113.232 | 316.29472 | |
| 883.96448 | | | 504.25984 | 365.36192 | 222.6896 | |
| 205.32736 | | | 95.11488 | | 196.2688 | |
| 126.81984 | | | 316.29472 | | 139.6528 | |
| 366.1168 | | | | | 111.72224 | |
| 443.11456 | | | | | 275.5312 | |
| 277.04096 | | | | | 206.83712 | |
| 114.74176 | | | | | 249.86528 | |
| 240.05184 | | | | | 144.93696 | |
| | | | | | 83.0368 | |
| | | | | | 277.79584 | |
| | | | | | 203.06272 | |
| | | | | | 167.58336 | |

| KI16 | KI17 | KI18 | KI19 | KI20 | KI21 | KI22 |
|---|---|---|---|---|---|---|
| 102.66368 | 130.59424 | 342.71552 | 313.2752 | 801.68256 | 186.45536 | 534.45504 |
| 983.60864 | 628.06016 | 226.464 | 110.21248 | 462.74144 | 649.95168 | 222.6896 |
| 284.58976 | 406.12544 | 763.93856 | 243.82624 | 466.51584 | 421.97792 | 145.69184 |
| 200.0432 | 260.4336 | 203.06272 | 141.91744 | 181.1712 | 336.67648 | 274.02144 |
| 160.03456 | 175.13216 | 879.4352 | 230.2384 | 348.75456 | 133.61376 | 281.57024 |
| 92.85024 | 143.4272 | 321.57888 | 252.8848 | 1120.2419 | 122.29056 | 175.88704 |
| 101.9088 | 258.92384 | 505.01472 | 366.87168 | 181.92608 | | 135.12352 |
| 117.0064 | 202.30784 | 643.91264 | 253.63968 | 286.8544 | | 168.33824 |
| 143.4272 | 92.09536 | 205.32736 | 170.60288 | 204.57248 | | |
| 124.5552 | 321.57888 | 126.06496 | 230.2384 | 1208.5629 | | |
| | 171.35776 | 153.99552 | 75.488 | 382.72416 | | |
| | 113.232 | 160.03456 | 153.99552 | 886.984 | | |
| | 72.46848 | 188.72 | 135.8784 | 890.00352 | | |
| | 220.42496 | 129.83936 | 358.568 | 834.1424 | | |
| | 315.53984 | 366.1168 | 146.44672 | 1281.0314 | | |
| | | 133.61376 | 747.3312 | 675.6176 | | |
| | | | 117.76128 | 202.30784 | | |
| | | | 104.17344 | 678.63712 | | |
| | | | 227.97376 | 1503.721 | | |
| | | | 283.83488 | 757.14464 | | |
| | | | 263.45312 | 887.73888 | | |
| | | | 936.0512 | 849.99488 | | |
| | | | 1171.5738 | 777.5264 | | |
| | | | 178.90656 | 366.1168 | | |
| | | | 110.96736 | 717.89088 | | |
| | | | | 1068.9101 | | |
| | | | | 1745.2826 | | |
| | | | | 1082.4979 | | |
| | | | | 897.55232 | | |
| | | | | 618.24672 | | |
| | | | | 748.08608 | | |
| | | | | 1068.9101 | | |
| | | | | 547.288 | | |
| | | | | 751.1056 | | |
| | | | | 1254.6106 | | |
| | | | | 436.32064 | | |
| | | | | 529.92576 | | |

| KI23 | KI25 | KI26 | KI27 | KI28 | KI30 | KI31 |
|---|---|---|---|---|---|---|
| 167.58336 | 125.31008 | 215.89568 | 372.15584 | 190.98464 | 349.50944 | 172.56557 |
| 170.60288 | 148.71136 | 375.93024 | 221.17984 | 113.98688 | 169.09312 | 230.08742 |
| 394.80224 | 119.27104 | 153.99552 | 240.05184 | 107.94784 | 227.97376 | 172.56557 |
| 818.28992 | | 189.47488 | 209.10176 | 834.89728 | 465.00608 | 134.21766 |
| 123.04544 | | 252.8848 | 163.80896 | 250.62016 | 454.43776 | 402.65299 |
| 495.95616 | | 103.41856 | 384.23392 | | 172.86752 | 479.3488 |
| 109.4576 | | 408.39008 | 293.64832 | | 409.89984 | |
| 137.38816 | | 168.33824 | 189.47488 | | 249.1104 | |
| 269.49216 | | 255.14944 | 99.64416 | | 412.91936 | |
| 228.72864 | | 161.54432 | 166.82848 | | 249.1104 | |
| 723.17504 | | 241.5616 | 94.36 | | 249.1104 | |
| 117.76128 | | 343.4704 | 191.73952 | | 563.89536 | |
| 594.09056 | | 291.38368 | | | 529.17088 | |
| 607.6784 | | 243.82624 | | | 415.184 | |
| 398.57664 | | | | | 200.79808 | |
| 472.55488 | | | | | 138.89792 | |
| 250.62016 | | | | | 560.12096 | |
| 363.09728 | | | | | | |
| 542.00384 | | | | | | |
| 635.60896 | | | | | | |
| 465.76096 | | | | | | |

| KI32 | KI33 | KI34 | KI35 | KI36 | KI37 | KI38 |
|---|---|---|---|---|---|---|
| 440.84992 | 283.83488 | 157.01504 | 150.976 | 345.73504 | 132.104 | 128.3296 |
| 160.03456 | 453.68288 | 247.60064 | 192.4944 | 185.70048 | 102.66368 | 284.58976 |
| 132.85888 | 171.35776 | 356.30336 | 178.90656 | 351.77408 | 124.5552 | 152.48576 |
| 114.74176 | 126.06496 | 99.64416 | 92.09536 | 1164.025 | 132.85888 | 114.74176 |
| 275.5312 | 274.77632 | 94.36 | 120.7808 | 276.28608 | 90.5856 | 436.32064 |
| 122.29056 | | 84.54656 | 154.7504 | 185.70048 | 98.1344 | 812.25088 |
| 104.92832 | | 122.29056 | 132.104 | 205.32736 | 84.54656 | 120.7808 |
| 121.53568 | | 187.21024 | 104.92832 | 135.8784 | | 116.25152 |
| 292.13856 | | 107.19296 | | 378.94976 | | 1224.4154 |
| 404.61568 | | 252.12992 | | 333.65696 | | 224.95424 |
| | | 102.66368 | | 77.75264 | | 677.88224 |
| | | 107.19296 | | 144.18208 | | 317.0496 |
| | | | | 633.34432 | | 520.8672 |
| | | | | 523.13184 | | |
| | | | | 129.08448 | | |
| | | | | 209.10176 | | |
| | | | | 190.22976 | | |
| | | | | 132.104 | | |
| | | | | 234.76768 | | |
| | | | | 497.46592 | | |
| | | | | 351.0192 | | |

Isolate B - schizont area (µm2)

| DMSO | KI1 | KI2 | KI3 | KI4 | KI5 | KI6 |
|---|---|---|---|---|---|---|
| 240.80672 | 1278.0118 | 1434.272 | 1138.359 | 1142.1334 | 277.04096 | 857.54368 |
| 668.0688 | 575.97344 | 1558.8272 | 742.04704 | 277.79584 | 759.40928 | 1231.9642 |
| 474.06464 | 919.44384 | 1237.2483 | 670.33344 | 399.33152 | 585.032 | 869.62176 |
| 425.75232 | 1475.0355 | 458.96704 | 634.0992 | 532.94528 | 2827.7805 | 1182.897 |
| 245.336 | 347.2448 | 897.55232 | 967.75616 | 1480.3197 | 485.38784 | 643.15776 |
| 428.77184 | 1081.743 | 1287.8253 | 2115.1738 | 508.03424 | 1742.263 | 357.81312 |
| 208.34688 | 185.70048 | 1778.4973 | 512.56352 | 415.184 | 1573.9248 | 691.47008 |
| 352.52896 | 1150.4371 | 1274.9923 | 420.46816 | 1157.231 | 689.20544 | 1002.4806 |
| 202.30784 | 578.23808 | 1521.8381 | 475.5744 | 274.02144 | 2423.9197 | 180.41632 |
| 578.99296 | 687.69568 | 1746.7923 | 797.90816 | 1073.4394 | 1621.4822 | 557.85632 |
| 542.00384 | 2797.5853 | 732.98848 | 1731.6947 | 236.27744 | 695.24448 | 1010.0294 |
| 341.96064 | 690.7152 | 646.17728 | 343.4704 | 559.36608 | 1878.8963 | 2170.28 |
| 674.86272 | 357.81312 | 770.73248 | 251.37504 | 2320.5011 | 817.53504 | 706.56768 |
| 753.37024 | 963.98176 | 619.75648 | 637.11872 | 2509.2211 | 1783.0266 | 991.91232 |
| 1234.2288 | 399.33152 | 557.10144 | 468.0256 | 279.3056 | 2100.831 | 548.04288 |
| 873.39616 | 468.78048 | 851.50464 | 1905.3171 | 144.18208 | 221.93472 | 1561.0918 |
| 1041.7344 | 1013.8038 | 1952.8746 | 979.07936 | 316.29472 | 391.02784 | 1345.1962 |
| 631.07968 | | 625.04064 | | 372.91072 | 1025.127 | 862.82784 |
| 298.93248 | | 1194.975 | | 636.36384 | | 2485.065 |
| 197.02368 | | 354.7936 | | 1090.0467 | | 1823.0352 |
| 542.00384 | | 1490.888 | | 338.18624 | | |
| 1706.0288 | | 1874.367 | | 670.33344 | | |
| 720.15552 | | 990.40256 | | 397.06688 | | |
| 258.92384 | | 648.44192 | | 445.3792 | | |
| 474.81952 | | 1955.1392 | | 194.00416 | | |
| 447.64384 | | 2057.8029 | | 2070.6358 | | |
| 421.22304 | | 519.35744 | | 1001.7258 | | |
| 1133.8298 | | 822.8192 | | | | |
| 1420.6842 | | 136.63328 | | | | |
| 883.2096 | | | | | | |
| 569.9344 | | | | | | |
| 1060.6064 | | | | | | |
| 840.93632 | | | | | | |
| 212.12128 | | | | | | |
| 418.20352 | | | | | | |
| 107.94784 | | | | | | |
| 619.0016 | | | | | | |
| 665.80416 | | | | | | |
| 1358.784 | | | | | | |
| 734.49824 | | | | | | |
| 847.73024 | | | | | | |

| |
|---|
| 1326.3242 |
| 1305.9424 |
| 2679.824 |
| 831.12288 |
| 1282.5411 |
| 416.69376 |
| 488.40736 |
| 628.81504 |
| 779.03616 |
| 553.32704 |
| 379.70464 |
| 1122.5066 |
| 1029.6563 |
| 446.13408 |
| 685.43104 |
| 1514.2893 |
| 2306.1584 |
| 1710.5581 |
| 2497.8979 |
| 2031.3821 |
| 1209.3178 |
| 1914.3757 |
| 290.6288 |
| 670.33344 |
| 874.15104 |
| 477.08416 |
| 1920.4147 |
| 634.85408 |
| 1069.665 |
| 886.22912 |
| 644.66752 |
| 1366.3328 |
| 243.82624 |
| 1560.337 |
| 454.43776 |
| 793.37888 |
| 341.20576 |
| 196.2688 |
| 1075.704 |
| 925.48288 |
| 305.7264 |
| 957.18784 |
| 374.42048 |

| |
|---|
| 468.0256 |
| 662.02976 |
| 666.55904 |
| 306.48128 |
| 719.40064 |
| 185.70048 |
| 380.45952 |
| 248.35552 |
| 455.19264 |
| 438.58528 |
| 446.13408 |
| 2287.2864 |
| 634.85408 |
| 436.32064 |

| KI7 | KI8 | KI10 | KI11 | KI13 | KI14 | KI15 |
|---|---|---|---|---|---|---|
| 1072.6845 | 1116.4675 | 452.17312 | 2066.1066 | 837.16192 | 1104.3894 | 810.74112 |
| 465.00608 | 730.72384 | 1647.903 | 684.67616 | 1650.9226 | 336.67648 | 178.90656 |
| 1303.6778 | 564.65024 | 1413.8902 | 1290.8448 | 610.69792 | 348.75456 | 546.53312 |
| 324.5984 | 691.47008 | 353.28384 | 616.73696 | 2236.7094 | 793.37888 | 705.8128 |
| 652.21632 | 673.35296 | 1773.968 | 172.86752 | 1000.9709 | 548.04288 | 1406.3414 |
| 453.68288 | 597.86496 | 726.19456 | 640.13824 | 2807.3987 | 563.89536 | 1820.0157 |
| 780.54592 | 1982.3149 | 468.0256 | 795.64352 | 1041.7344 | 265.71776 | 252.8848 |
| 1786.801 | 1177.6128 | 705.05792 | 188.72 | 670.33344 | 1038.7149 | 571.44416 |
| 1035.6954 | 2482.0454 | 1326.3242 | 571.44416 | 436.32064 | 1212.3373 | 1776.2326 |
| 663.53952 | 1651.6774 | 344.22528 | 867.35712 | 799.41792 | 1194.2202 | 485.38784 |
| 2026.8528 | 2623.208 | 989.64768 | 2670.0106 | 1881.9158 | 1434.272 | 1668.2848 |
| 3431.6845 | 2152.1629 | 1503.721 | 1276.5021 | 870.37664 | 674.10784 | 934.54144 |
| 1008.5197 | 883.2096 | 1895.5037 | 1767.929 | 1709.0483 | 927.74752 | 1416.1549 |
| 418.20352 | 651.46144 | 1379.1658 | 2017.0394 | 1147.4176 | 277.79584 | 1093.0662 |
| 1119.487 | 866.60224 | 2516.015 | 295.91296 | 1074.9491 | 1940.7965 | 890.00352 |
| 452.17312 | 164.56384 | 1813.2218 | 486.8976 | 298.1776 | 1112.6931 | 249.86528 |
| 1087.7821 | 615.2272 | 1455.4086 | 906.61088 | 691.47008 | 449.90848 | 2536.3968 |
| 1001.7258 | 628.06016 | 389.51808 | 887.73888 | 713.3616 | 1604.8749 | |
| | 1331.6083 | 582.76736 | 1292.3546 | | 717.89088 | |
| | 621.26624 | 842.44608 | 458.21216 | | 1228.1898 | |
| | 2237.4643 | 736.76288 | 736.008 | | 345.73504 | |
| | | 954.16832 | 579.74784 | | 192.4944 | |
| | | 1546.7491 | 619.75648 | | 1098.3504 | |
| | | 1726.4106 | 523.13184 | | 132.85888 | |
| | | 355.54848 | 1016.0685 | | 1376.9011 | |
| | | 589.56128 | 1295.3741 | | 725.43968 | |
| | | | 453.68288 | | 2754.5571 | |
| | | | 3493.5846 | | 393.29248 | |
| | | | | | 769.9776 | |
| | | | | | 1266.6886 | |

| KI16 | KI17 | KI18 | KI19 | KI20 | KI21 | KI22 |
|---|---|---|---|---|---|---|
| 163.80896 | 1629.031 | 440.84992 | 667.31392 | 2053.2736 | 1749.057 | 763.93856 |
| 745.82144 | 655.23584 | 1533.9162 | 1584.4931 | 2304.6486 | 1130.8102 | 410.65472 |
| 516.33792 | 430.2816 | 270.24704 | 1933.2477 | 3096.5178 | 971.53056 | 2255.5814 |
| 346.48992 | 428.01696 | 2217.8374 | 838.67168 | 1892.4842 | 1873.6122 | 745.82144 |
| 824.32896 | 847.73024 | 2227.6509 | 279.3056 | 3100.2922 | 493.69152 | 1043.999 |
| 1755.8509 | 634.85408 | 2134.8006 | 160.03456 | 3034.6176 | 451.41824 | 580.50272 |
| 732.2336 | 1342.9315 | 1083.2528 | 1997.4125 | 1636.5798 | 1203.2787 | 1024.3722 |
| 194.75904 | 913.4048 | 563.89536 | 2324.2755 | 2705.4899 | 697.50912 | 1238.7581 |
| 1107.409 | 554.8368 | 360.83264 | | 1915.8854 | 1419.9293 | 271.7568 |
| 953.41344 | 1921.1696 | 2029.1174 | | 2035.9114 | 695.24448 | 479.3488 |
| 1489.3782 | 383.47904 | 783.56544 | | 2592.2579 | 3452.0662 | 1445.5952 |
| | 1340.6669 | 660.52 | | 1439.5562 | 1269.7082 | 601.63936 |
| | 951.90368 | 826.5936 | | 3251.2682 | | 1416.9098 |
| | 1210.8275 | | | 3941.2285 | | 1583.7382 |
| | 167.58336 | | | 1515.799 | | 937.56096 |
| | 1827.5645 | | | 2233.6899 | | 630.3248 |
| | 1145.9078 | | | 1516.5539 | | 553.32704 |
| | 1105.8992 | | | 918.68896 | | 782.05568 |
| | 313.2752 | | | 336.67648 | | 934.54144 |
| | 2265.3949 | | | 1761.135 | | 1453.8989 |
| | 319.31424 | | | 885.47424 | | 766.2032 |
| | 1782.2717 | | | 713.3616 | | |
| | 314.03008 | | | 1639.5994 | | |
| | 1439.5562 | | | 1090.8016 | | |
| | 1207.808 | | | 1687.9117 | | |
| | 873.39616 | | | 3350.1574 | | |
| | 479.3488 | | | 1262.1594 | | |
| | 1165.5347 | | | 3147.0947 | | |
| | 791.11424 | | | 1185.9165 | | |
| | 1730.9398 | | | 1805.673 | | |
| | 2022.3235 | | | 763.18368 | | |
| | 1325.5693 | | | 4320.9331 | | |
| | 503.50496 | | | 2054.0285 | | |
| | 1733.9594 | | | 1391.2438 | | |
| | 412.91936 | | | 2212.5533 | | |
| | 697.50912 | | | 3285.9926 | | |
| | 593.33568 | | | | | |
| | 1610.9139 | | | | | |
| | 1350.4803 | | | | | |
| | 1186.6714 | | | | | |
| | 1108.9187 | | | | | |

| |
|---|
| 293.64832 |
| 371.40096 |
| 2009.4906 |
| 751.1056 |
| 846.97536 |
| 466.51584 |

| KI23 | KI25 | KI26 | KI27 | KI28 | KI30 | KI31 |
|---|---|---|---|---|---|---|
| 415.184 | 194.00416 | 911.14016 | 246.09088 | 500.48544 | 551.81728 | 239.29696 |
| 954.9232 | 677.88224 | 561.63072 | 351.0192 | 443.86944 | 295.15808 | 752.61536 |
| 309.5008 | 599.37472 | 617.49184 | 694.4896 | 1162.5152 | 311.01056 | 157.01504 |
| 578.23808 | 212.12128 | 980.58912 | 726.19456 | 595.60032 | 1650.9226 | 594.84544 |
| 572.19904 | 830.368 | 643.15776 | 846.22048 | 1700.7446 | 1065.1357 | 128.3296 |
| 1713.5776 | 247.60064 | 517.84768 | 799.41792 | 423.48768 | 249.86528 | 264.208 |
| 1183.6518 | 314.78496 | 463.49632 | 643.15776 | 783.56544 | 1077.2138 | 126.06496 |
| 1047.0186 | 1006.255 | 479.3488 | 1136.0944 | 1308.9619 | 1147.4176 | 136.63328 |
| 1994.393 | 1143.6432 | 1072.6845 | 1478.8099 | 1064.3808 | 531.43552 | 171.35776 |
| 2131.0262 | 685.43104 | 1370.8621 | 196.2688 | 1618.4627 | 303.46176 | 178.15168 |
| 1127.7907 | 1547.504 | 1327.8339 | 589.56128 | 577.4832 | 184.19072 | 159.27968 |
| 2370.3232 | 534.45504 | 2252.5619 | 384.23392 | 1030.4112 | 280.06048 | 254.39456 |
| 323.84352 | | 2401.2733 | 347.99968 | 308.74592 | 684.67616 | 695.99936 |
| 754.12512 | | 1655.4518 | 137.38816 | 410.65472 | 1382.9402 | 560.12096 |
| 1552.0333 | | 1610.159 | 300.44224 | 1037.2051 | 678.63712 | 338.18624 |
| 696.75424 | | 625.04064 | 415.93888 | 324.5984 | 1231.9642 | 470.29024 |
| 1379.9206 | | 1512.7795 | 780.54592 | 191.73952 | 834.1424 | 166.82848 |
| 1140.6237 | | 1545.9942 | 210.61152 | 314.78496 | 895.28768 | 640.13824 |
| 2009.4906 | | 2293.3254 | 110.96736 | 631.83456 | 501.9952 | 914.91456 |
| | | 2086.4883 | 470.29024 | 474.81952 | 941.33536 | 355.54848 |
| | | 458.21216 | 292.89344 | 392.5376 | 249.1104 | |
| | | 1339.912 | 162.2992 | 413.67424 | 99.64416 | |
| | | 1380.6755 | | 168.33824 | 483.87808 | |
| | | 2203.4947 | | 781.3008 | | |
| | | 1626.7664 | | | | |
| | | 2862.505 | | | | |
| | | 1500.7014 | | | | |
| | | 1715.8422 | | | | |
| | | 1521.8381 | | | | |
| | | 122.29056 | | | | |

| KI32 | KI33 | KI34 | KI35 | KI36 | KI37 | KI38 |
|---|---|---|---|---|---|---|
| 844.71072 | 1291.5997 | 1028.9014 | 846.22048 | 1228.1898 | 476.32928 | 1312.7363 |
| 118.51616 | 338.18624 | 896.04256 | 1839.6426 | 245.336 | 751.1056 | 158.5248 |
| 574.46368 | 947.3744 | 802.43744 | 1450.1245 | 1031.921 | 181.92608 | 665.80416 |
| 813.00576 | 2040.4406 | 1366.3328 | 905.856 | 360.83264 | 1494.6624 | 602.39424 |
| 775.26176 | 197.77856 | 1000.216 | 160.03456 | 490.672 | 280.81536 | 988.13792 |
| 628.81504 | 265.71776 | 785.83008 | 816.02528 | 194.75904 | 927.74752 | 744.31168 |
| 463.49632 | 598.61984 | 1800.3888 | 561.63072 | 618.24672 | 840.93632 | 914.15968 |
| 666.55904 | 372.15584 | 962.472 | 1081.743 | 881.69984 | 828.85824 | 723.92992 |
| 765.44832 | 372.91072 | 1696.2154 | 576.72832 | 470.29024 | 199.28832 | 463.49632 |
| 309.5008 | 318.55936 | 594.84544 | 942.09024 | 575.21856 | 462.74144 | 202.30784 |
| 600.1296 | 655.23584 | 311.76544 | 493.69152 | 691.47008 | 672.59808 | 638.62848 |
| 833.38752 | 1200.2592 | 919.44384 | 496.71104 | 907.36576 | 548.04288 | 223.44448 |
| 1210.8275 | 492.18176 | 990.40256 | 726.94944 | 1561.8467 | 602.39424 | 462.74144 |
| 729.21408 | 219.67008 | 883.96448 | 553.32704 | 637.8736 | 210.61152 | 575.21856 |
| 806.21184 | 438.58528 | 341.96064 | 400.0864 | 492.18176 | 1564.8662 | 390.27296 |
| 802.43744 | 325.35328 | 560.12096 | 271.7568 | 790.35936 | | 1724.1459 |
| 408.39008 | 1281.0314 | 1604.8749 | 378.94976 | 372.15584 | | 875.6608 |
| 1101.3699 | 2257.8461 | 189.47488 | 708.83232 | 442.35968 | | 972.28544 |
| 445.3792 | 679.392 | 1282.5411 | 344.98016 | 458.96704 | | 575.21856 |
| 619.75648 | 1567.1309 | | 316.29472 | 544.26848 | | 730.72384 |
| 250.62016 | 338.18624 | | 432.54624 | 223.44448 | | 318.55936 |
| 776.77152 | 195.51392 | | 233.25792 | 555.59168 | | 223.44448 |
| 232.50304 | 334.41184 | | 1063.6259 | | | 718.64576 |
| 208.34688 | | | 175.13216 | | | 999.46112 |
| 541.24896 | | | 945.86464 | | | 1151.192 |
| 279.3056 | | | 1330.0986 | | | 888.49376 |
| 1062.1162 | | | 192.4944 | | | |
| 1164.7798 | | | 502.75008 | | | |
| 167.58336 | | | 869.62176 | | | |
| 826.5936 | | | 159.27968 | | | |
| 1271.2179 | | | 464.2512 | | | |
| 2006.471 | | | 1680.3629 | | | |
| 1454.6538 | | | 1347.4608 | | | |
| 1308.9619 | | | 465.00608 | | | |
| 1209.3178 | | | 1269.7082 | | | |

Isolate C - schizont area (µm2)

| DMSO | KI1 | KI2 | KI3 | KI4 | KI5 | KI6 |
|---|---|---|---|---|---|---|
| 1080.2333 | 612.96256 | 289.87392 | 2133.2909 | 511.05376 | 2495.6333 | 1779.2522 |
| 454.43776 | 1326.3242 | 875.6608 | 328.3728 | 410.65472 | 1444.8403 | 1255.3654 |
| 2229.9155 | 1813.9766 | 736.76288 | 474.81952 | 271.7568 | 565.40512 | 2035.9114 |
| 667.31392 | 474.81952 | | 856.7888 | 393.29248 | 1391.2438 | 980.58912 |
| 502.75008 | 155.50528 | | 912.64992 | 254.39456 | 784.32032 | 465.76096 |
| 831.87776 | 994.17696 | | 270.24704 | 467.27072 | 2022.3235 | 818.28992 |
| 970.0208 | 1053.0576 | | 1166.2896 | 300.44224 | 893.02304 | 309.5008 |
| 584.27712 | 152.48576 | | | | 546.53312 | 1396.528 |
| 670.33344 | 357.05824 | | | | 1012.2941 | 1937.0221 |
| 831.87776 | 856.7888 | | | | 948.88416 | 397.06688 |
| 2206.5142 | 542.00384 | | | | 726.94944 | 1795.8595 |
| 177.3968 | | | | | 762.4288 | 1011.5392 |
| 886.984 | | | | | | |
| 1397.2829 | | | | | | |
| 1897.0134 | | | | | | |
| 1413.1354 | | | | | | |
| 431.03648 | | | | | | |
| 178.90656 | | | | | | |
| 308.74592 | | | | | | |
| 1826.0547 | | | | | | |
| 1652.4323 | | | | | | |
| 674.10784 | | | | | | |
| 655.99072 | | | | | | |
| 1442.5757 | | | | | | |
| 1468.9965 | | | | | | |
| 1394.2634 | | | | | | |
| 1864.5536 | | | | | | |

| KI7 | KI8 | KI10 | KI11 | KI13 | KI14 | KI15 |
|---|---|---|---|---|---|---|
| 782.81056 | 992.6672 | 455.94752 | 671.8432 | 1768.6838 | 1193.4653 | 1870.5926 |
| 1217.6214 | 1972.5014 | 163.05408 | 820.55456 | 1090.8016 | 722.42016 | 1949.855 |
| 572.95392 | 264.96288 | 1437.2915 | 434.81088 | 1768.6838 | 197.02368 | 1281.7862 |
| 695.99936 | 424.99744 | 1706.7837 | 952.65856 | 1404.8317 | 713.3616 | 1855.495 |
| 220.42496 | 890.00352 | 1595.0614 | 1264.424 | 198.53344 | 285.34464 | 932.2768 |
| 264.208 | 1367.8426 | 581.2576 | 881.69984 | 461.98656 | 704.30304 | 871.13152 |
| | 1182.897 | 1201.0141 | 690.7152 | 860.5632 | 1729.4301 | |
| | 1703.0093 | 399.33152 | | 935.29632 | 1284.0509 | |
| | 615.98208 | | | 1126.281 | 340.45088 | |
| | 419.71328 | | | 984.36352 | | |
| | 223.44448 | | | 608.43328 | | |
| | 1050.793 | | | 666.55904 | | |
| | | | | 786.58496 | | |
| | | | | 1841.1523 | | |

| KI16 | KI17 | KI18 | KI19 | KI20 | KI21 | KI22 |
|---|---|---|---|---|---|---|
| 763.18368 | 1464.4672 | 987.38304 | 631.83456 | 621.26624 | 534.45504 | 1297.6387 |
| 753.37024 | 1856.2499 | 508.03424 | 1820.7706 | 3067.0774 | 1633.5603 | 1391.2438 |
| 1298.3936 | 1650.9226 | 625.04064 | 1000.9709 | 2796.8304 | 424.24256 | 320.06912 |
| 423.48768 | 391.78272 | 891.51328 | 1900.033 | 380.45952 | 901.32672 | 548.04288 |
| 914.15968 | 181.1712 | 516.33792 | 1786.0461 | 1564.1114 | 978.32448 | 1542.9747 |
| 766.95808 | 2053.2736 | | 1746.7923 | 3585.68 | 268.73728 | 281.57024 |
| 499.73056 | 219.67008 | | 1293.8643 | 295.15808 | 1589.7773 | 1194.975 |
| 946.61952 | | | 1285.5606 | 1890.2195 | 363.85216 | |
| 1267.4435 | | | 2028.3626 | 1100.615 | 885.47424 | |
| | | | 1252.3459 | 2793.056 | 355.54848 | |
| | | | | 2019.304 | 1936.2672 | |
| | | | | 3763.8317 | 1558.0723 | |
| | | | | 2482.0454 | 1358.784 | |

| KI23 | KI25 | KI26 | KI27 | KI28 | KI30 | KI31 |
|---|---|---|---|---|---|---|
| 1154.2115 | 529.92576 | 3394.6954 | 951.90368 | 402.35104 | 625.04064 | 356.30336 |
| 1139.1139 | 582.76736 | 1408.6061 | 149.46624 | 2211.7984 | 394.80224 | 1296.129 |
| 1087.0272 | 1256.8752 | 1272.7277 | 260.4336 | 883.96448 | 260.4336 | |
| 677.12736 | 305.7264 | 903.59136 | 869.62176 | 1274.9923 | 449.1536 | |
| 1139.1139 | 1065.8906 | 695.99936 | 212.12128 | 211.3664 | 311.76544 | |
| 1009.2746 | 1191.9555 | 832.63264 | 432.54624 | 123.80032 | 550.30752 | |
| 778.28128 | 462.74144 | 795.64352 | 1805.673 | 489.91712 | 328.3728 | |
| 813.76064 | 861.31808 | 859.80832 | 772.24224 | 713.3616 | 128.3296 | |
| 1013.049 | 181.1712 | 2214.8179 | 657.50048 | 2595.2774 | 158.5248 | |
| 1900.033 | 147.2016 | 2020.8138 | 175.88704 | | 505.01472 | |
| 762.4288 | 526.15136 | 322.33376 | 1355.7645 | | 885.47424 | |
| 1491.6429 | 452.17312 | 2512.9955 | 1010.7843 | | 1729.4301 | |
| 183.43584 | 1053.8125 | 550.30752 | 1677.3434 | | | |
| 1260.6496 | 899.06208 | 1057.5869 | 629.56992 | | | |
| | 1182.897 | 1902.2976 | 571.44416 | | | |
| | | 2536.3968 | 1682.6275 | | | |
| | | 246.09088 | | | | |
| | | 1972.5014 | | | | |

| KI32 | KI33 | KI34 | KI35 | KI36 | KI37 | KI38 |
|---|---|---|---|---|---|---|
| 933.03168 | 737.51776 | 411.4096 | 1743.7728 | 708.83232 | 542.75872 | 1378.4109 |
| 525.39648 | 508.03424 | 1265.9338 | 2124.2323 | 1056.832 | 483.1232 | 1139.8688 |
| 414.42912 | 148.71136 | 1266.6886 | 201.55296 | 689.20544 | 847.73024 | 568.42464 |
| 175.13216 | 1182.897 | 988.8928 | 1521.0832 | 1093.8211 | 729.96896 | 928.5024 |
| 468.0256 | 519.35744 | 2041.9504 | 1881.9158 | 982.09888 | 982.09888 | 585.032 |
| 502.75008 | 432.54624 | 1764.9094 | 711.85184 | 227.97376 | 684.67616 | 824.32896 |
| 246.84576 | 455.94752 | 1339.1571 | 116.25152 | 1002.4806 | 301.952 | 481.61344 |
| 778.28128 | 1022.1075 | 342.71552 | 606.92352 | 244.58112 | 2320.5011 | 641.648 |
| | 88.32096 | 1185.1616 | 1079.4784 | 2246.5229 | 1536.9357 | 498.97568 |
| | 913.4048 | 960.20736 | 742.04704 | 795.64352 | 173.6224 | 258.92384 |
| | 1249.3264 | 1536.9357 | | | 572.95392 | 1425.2134 |
| | | 325.35328 | | | 322.33376 | 588.05152 |
| | | 1096.8406 | | | 388.7632 | 292.13856 |
| | | 2046.4797 | | | 123.80032 | 294.4032 |
| | | 1265.9338 | | | 862.82784 | |

**Nucleus counts**

isolate A

| DMSO | KI1 | KI2 | KI3 | KI4 | KI5 | KI6 |
|---|---|---|---|---|---|---|
| 4668 | 5086 | 4945 | 5023 | 4936 | 5235 | 4608 |
| 5227 | 5093 | 5325 | 5100 | 4840 | 5101 | 5258 |
| 5372 | 5244 | 4900 | 5361 | 5149 | 4995 | 5441 |

isolate B

| DMSO | KI1 | KI2 | KI3 | KI4 | KI5 | KI6 |
|---|---|---|---|---|---|---|
| 7614 | 5275 | 6717 | 5797 | 6639 | 4840 | 7010 |
| 5283 | 6242 | 7208 | 7546 | 6262 | 6282 | 5399 |
| 7691 | 6117 | 6275 | 6607 | 6253 | 5411 | 5620 |
| 6366 | | | | | | |
| 6091 | | | | | | |
| 6584 | | | | | | |
| 6256 | | | | | | |
| 6562 | | | | | | |
| 6833 | | | | | | |

isolate C

| DMSO | KI1 | KI2 | KI3 | KI4 | KI5 | KI6 |
|---|---|---|---|---|---|---|
| 5968 | 6791 | 6348 | 6113 | 7321 | 6512 | 6894 |
| 7013 | 6701 | 6672 | 4470 | 6112 | 6183 | 7803 |
| 5219 | 6966 | 6383 | 7197 | 6675 | 6589 | 5492 |
| 6409 | | | | | | |
| 7200 | | | | | | |
| 7028 | | | | | | |
| 6686 | | | | | | |
| 6566 | | | | | | |
| 6653 | | | | | | |

| KI7 | KI8 | KI10 | KI11 | KI13 | KI14 | KI15 |
|---|---|---|---|---|---|---|
| 5070 | 5150 | 4721 | 5370 | 4509 | 5030 | 5061 |
| 5335 | 5427 | 5131 | 5539 | 5180 | 5367 | 5183 |
| 5481 | 4938 | 5511 | 5412 | 5213 | 4811 | 5462 |

| KI7 | KI8 | KI10 | KI11 | KI13 | KI14 | KI15 |
|---|---|---|---|---|---|---|
| 5205 | 6191 | 6842 | 4768 | 6223 | 6989 | 6181 |
| 6160 | 6277 | 6018 | 5236 | 6740 | 6126 | 6242 |
| 6319 | 5790 | 6448 | 5848 | 6930 | 6128 | 6652 |

| KI7 | KI8 | KI10 | KI11 | KI13 | KI14 | KI15 |
|---|---|---|---|---|---|---|
| 5739 | 6445 | 6267 | 7084 | 6697 | 6132 | 6551 |
| 7113 | 6732 | 4626 | 6796 | 5999 | 7661 | 6847 |
| 6599 | 6889 | 5393 | 5976 | 6670 | 6036 | 6663 |

| KI16 | KI17 | KI18 | KI19 | KI20 | KI21 | KI22 |
|---|---|---|---|---|---|---|
| 5223 | 5254 | 4620 | 5063 | 4857 | 4960 | 5107 |
| 4978 | 5309 | 4766 | 4794 | 5027 | 5069 | 5089 |
| 5376 | 5243 | 4981 | 4962 | 4590 | 4882 | 5032 |

| KI16 | KI17 | KI18 | KI19 | KI20 | KI21 | KI22 |
|---|---|---|---|---|---|---|
| 6349 | 5859 | 6198 | 4697 | 6663 | 6213 | 6283 |
| 5700 | 6405 | 4873 | 5835 | 6606 | 6174 | 6473 |
| 6048 | 6371 | 6100 | 6542 | 5890 | 5431 | 6623 |

| KI16 | KI17 | KI18 | KI19 | KI20 | KI21 | KI22 |
|---|---|---|---|---|---|---|
| 6470 | 6259 | 6573 | 6395 | 6226 | 6524 | 6693 |
| 6534 | 4532 | 6335 | 5775 | 7326 | 6305 | 4635 |
| 6117 | 5927 | 6852 | 6364 | 6092 | 6485 | 6160 |

| KI23 | KI25 | KI26 | KI27 | KI28 | KI29 | KI30 |
| --- | --- | --- | --- | --- | --- | --- |
| 4468 | 3813 | 5048 | 5015 | 4740 | 1452 | 4606 |
| 4671 | 4210 | 5281 | 4668 | 5361 | 1620 | 4818 |
| 4872 | 4153 | 4952 | 4935 | 5087 | 1953 | 4758 |

| KI23 | KI25 | KI26 | KI27 | KI28 | KI29 | KI30 |
| --- | --- | --- | --- | --- | --- | --- |
| 6674 | 5188 | 7488 | 6495 | 6007 | 5204 | 6392 |
| 5723 | 5963 | 6565 | 6432 | 6121 | 5435 | 5594 |
| 6013 | 5707 | 6675 | 6667 | 6050 | 2918 | 4956 |

| KI23 | KI25 | KI26 | KI27 | KI28 | KI29 | KI30 |
| --- | --- | --- | --- | --- | --- | --- |
| 6663 | 5928 | 6872 | 6862 | 6466 | 4623 | 6298 |
| 6332 | 6187 | 7087 | 6752 | 6700 | 2961 | 5878 |
| 6599 | 5937 | 6841 | 6463 | 6708 | 3310 | 6233 |

| KI31 | KI32 | KI33 | KI34 | KI35 | KI36 | KI37 |
|---|---|---|---|---|---|---|
| 4584 | 5096 | 5078 | 5172 | 4791 | 4725 | 4801 |
| 4897 | 4990 | 4858 | 5028 | 5157 | 4912 | 5362 |
| 5124 | 4800 | 4972 | 5810 | 5271 | 5058 | 4539 |

| KI31 | KI32 | KI33 | KI34 | KI35 | KI36 | KI37 |
|---|---|---|---|---|---|---|
| 3911 | 6852 | 6950 | 4693 | 6099 | 6666 | 4883 |
| 6473 | 6896 | 6512 | 2379 | 7195 | 6074 | 5616 |
| 5976 | 6506 | 7414 | 4765 | 5710 | 6203 | 6055 |

| KI31 | KI32 | KI33 | KI34 | KI35 | KI36 | KI37 |
|---|---|---|---|---|---|---|
| 6632 | 6839 | 6973 | 6658 | 6846 | 7112 | 6714 |
| 6167 | 7869 | 6977 | 6326 | 5122 | 6577 | 6825 |
| 6876 | 6467 | 6452 | 6639 | 6620 | 6442 | 6222 |

KI38

4928
1959
4282

KI38

6077
6094
5538

KI38

6291
6546
5914

schizonts/well

| | KI31 - SU11274 | | |
|---|---|---|---|
| NT (DMSO) | 41 | 44 | 28 |
| 100nM | 36 | 33 | 27 |
| 500nM | 21 | 12 | 16 |
| 1uM | 8 | 4 | 2 |
| 10uM | 1 | 0 | 0 |

| | KI6 - casein kinae inhib D44 | | |
|---|---|---|---|
| NT (DMSO) | 33 | 32 | 32 |
| 100nM | 39 | 26 | 24 |
| 500nM | 27 | 28 | 0 |
| 1uM | 54 | 31 | 27 |
| 10uM | 3 | 1 | 11 |
